# Environmentally induced variation in sperm sRNAs is linked to gene expression and transposable elements in zebrafish offspring

**DOI:** 10.1101/2024.12.04.626822

**Authors:** Alice M. Godden, Willian T.A.F. Silva, Berrit Kiehl, Cécile Jolly, Leighton Folkes, Ghazal Alavioon, Simone Immler

## Abstract

Environmental factors affect not only paternal condition but may translate into the following generations where sperm-mediated small RNAs (sRNAs) can contribute to the transmission of paternal effects. SRNAs play a key role in the male germ line in genome maintenance and repair, and particularly in response to environmental stress and the resulting increase in transposable element (TE) activity. Here, we investigated how the social environment (high competition, low competition) of male zebrafish *Danio rerio* affects small RNAs in sperm and how these are linked to gene expression and TE activity in their offspring. In a first experiment, we collected sperm samples after exposing males to each social environment for two weeks to test for differentially expressed sperm micro-(miRNA) and piwi-interacting RNAs (piRNA). In a separate experiment, we performed *in vitro* fertilisations after one two-week period using a split-clutch design to control for maternal effects and collected embryos at 24 hours to test for differentially expressed genes and transposable elements. We developed new computational prediction tools to link sperm sRNAs with differentially expressed TEs and genes in the embryos. Our results support the idea that the molecular stress response in the male germ line has significant down-stream effects on the molecular pathways, and we provide a direct link between sRNAs, TEs and gene expression.

**Author summary:** The discovery that sperm transmit more than just the father’s genome to the next generation is relatively recent and the potential implications are far reaching. What we do not know is whether these non-genetic components contained in sperm play a role in adaptation to changing environments or whether they are simply a result of the father’s stress response that also affects the germ cells. Environmentally induced stress is known to trigger a response to maintain and repair the germ cells to ensure the production of high quality sperm and key factors in this response are small RNAS. Small RNAs are ‘guardians of the germ line’ and protect the germline genome against the activity of selfish genetic elements. We investigated how the social environment in male zebrafish affects the expression level of small RNAs in sperm and how these changes in small RNAs are linked to changes in the gene epxression and the activity of selfish genetic elements in the offspring. We developed specialised bioinformatic pipelines to provide a clear link between the response in the father’s germline to changes in the environment and gene expresssion in their offspring.

## Introduction

The importance of non-genetic inheritance of environmentally induced variation in paternal condition for offspring fitness is increasingly accepted (Bonduriansky *et al*, 2012; Bonduriansky and Day, 2009). A range of factors affect not only the condition of the father but also the resulting offspring including temperature (Salinas *et al*, 2013; Salinas and Munch, 2012), diet (Dunn and Bale, 2009; Huypens *et al*, 2016; Valtonen *et al*, 2012), social interactions (Saavedra-Rodriguez and Feig, 2013), and stress (Franklin *et al*, 2010; Kan *et al*, 2016; Weiss *et al*, 2011; Zaidan *et al*, 2013). While it is generally accepted that these effects may be transferred through epigenetic (namely, non-DNA encoded) factors such as methylation patterns (Jiang *et al*, 2013; Potok *et al*, 2013), chromatin structure (Lindeman *et al*, 2011) and small RNAs (sRNAs), (Gapp *et al*, 2014; Houri-Zeevi *et al*, 2020; Rechavi *et al*, 2014), it is unclear whether these factors ending up in sperm are adaptive or part of the repair and maintenance response of the male germ line to envrionmental stress (Godden and Immler, 2023). Here we focused on the possible role of sperm sRNAs in the transmission of inter-generational paternal effects and their link to transposable element (TE) activity and gene expression in the resulting offspring.

Among the sperm-mediated sRNAs, micro-(mi-) and piwi-interacting (pi-) RNAs are known to exhibit differential expression in sperm and may contribute to non-genetic inheritance (Immler, 2018; Rando, 2016). In adult male Norway rats *Rattus norwegicus* for example, heat stress altered gene expression as well as miRNA expression in spermatocytes and spermatids (Yadav *et al*, 2018), and in zebrafish *Danio rerio*, thermal stress affected the miRNA profile in testes (van Gelderen *et al*, 2022). MiRNA and piRNA profiles in sperm of male house mice *Mus musculus* exposed to trauma as juveniles differed from the profiles in sperm fo control mice, and these differences in sRNA expression were directly linked to changes in the behaviour of the resulting offspring (Gapp *et al*, 2020). Finally, long-term exposure to different sex ratios and diets in selection lines of the fruitfly *Drosophila melanogaster* led to differences in sperm miRNA profiles between the different lines (Hotzy *et al*, 2021).

However, sRNAs also have a key role in gene regulation and DNA maintenance and repair in the animal germ line (Bartel, 2004; Godden and Immler, 2023; Houwing *et al*, 2007). MiRNAs post-transcriptionally regulate gene expression through complementary seed region binding of the mature miRNA and the 3’ untranslated region of a mRNA; one miRNA can regulate many hundreds of genes, and as a result can fine-tune gene expression (Agarwal *et al*, 2015; Bartel, 2004; Thatcher *et al*, 2008). PiRNAs are involved in protecting the genome, particularly the germline genome from invasive TEs (Chang *et al*, 2022; Houwing *et al*, 2007). DNA transposons are the most abundant TEs in the zebrafish genome and contribute to the epigenetic state of the genome (Chang *et al*, 2022). Many DNA transposons exhibit testes-specific expression in zebrafish suggesting they may have tissue-specific regulatory functions (Lee *et al*, 2022). Long Terminal Repeat (LTR) elements are enriched very early in zebrafish embryo development during post-zygotic genome activation, and DNA TEs are expressed in later development (Chang *et al*, 2022). The enrichment for piRNAs complementary to LTR elements at these early stages of embryo development may therefore offer protection to the developing zebrafish germline in changing environments (Chang *et al*, 2022; Houwing *et al*, 2007).

While environmental factors such as temperature, diet and toxins are directly interacting with physiological processes (Barrionuevo and Burggren, 1999; Ton *et al*, 2003), the effects of the social environment on individuals is more complex. A common response to competitive social interactions is stress, which can potentially cause diseases in the individuals directly exposed to it (Garratt *et al*, 2016; McEwen and Stellar, 1993) and can affect the next generation (Franklin *et al*, 2010; Kan *et al*, 2016; Weiss *et al*, 2011; Zaidan *et al*, 2013). In fact, stressful environments lead to higher cortisol levels and in extreme cases death in subordinate zebrafish, as zebrafish act agressively and will establish dominance if kept in pairs (Dahlbom *et al*, 2011). As a consequence, social interactions and social hierarchy among males during competition for access to females are known to affect ejaculate and sperm traits in a wide variety of species (Fitzpatrick and Lupold, 2014), and dominant males produce more sperm compared to subordinate males (Filby *et al*, 2010). Stress experienced by males may also carry over into the following generations, potentially through the transmission of sRNAs in sperm. In male zebrafish for example, stress induced by an alarm cue prior to mating led to differential expression in sRNA profiles in sperm, where 12 out of 213 mature miRNAs, six out of a 569 piRNA clusters (all enriched), and 12 out of 55 tsRNA clusters showed differential expression (Ord *et al*, 2020). In addition, offspring of these stressed males exhibited altered stress responses, and increased thigmotaxis. We previously showed that the intensity of male-male competition experienced by fathers prior to siring offspring affects offspring performance, with males under high competition siring faster hatching offspring with lower survival rates (Zajitschek *et al*, 2014). Similarly, the social status of zebrafish males immediately before fertilisation affected offspring activity, and changes in social hierarchy had the strongest effects where males shifting from dominant to subordinate seemed to be experiencing the biggest change (Zajitschek *et al*, 2017). The mechanisms of these observed effects on offsping phenotype are currently not known.

The aim of our study was to understand how variation in the paternal social environment may affect sRNA profiles in sperm and whether these are linked to gene expression and TE activity during early embryo development in the resulting offspring. We exposed male zebrafish to an experimental setup where males were either kept in company of one male and one female (high competition) or in the company of two females (low competition). In a first experiment, we exposed each male to both treatments in random order for two weeks each and collected a total of two sperm samples, one at the end of each period. We conducted sRNA sequencing with a focus on mi-and piRNAs on the sperm samples and mRNA sequencing on the embryos to identify differentially expressed genes and TEs. In a second experiment, we exposed males to one of the two treatments for two weeks only, and performed *in vitro* fertilisations (IVF) at the end using a split-clutch design to control for maternal effects. We collected embryos at 24 hours post-fertilisation, a key stage in early embryo development with highly synchronised expression patterns (Mathavan *et al*, 2005), for whole transcriptome analyses. Subsequently, we were able to link differentially expressed sRNAs in sperm with differential expression of putative target genes and TEs in the offspring using newly developped bioinformatic tools.

## Materials and methods

All fish used in the experiment were adult male and female zebrafish from the AB wild-type strain obtained from ZIRC and bred under a carefully designed outbreeding regime for one generation in the SciLifeLab zebrafish facility at Uppsala University. The fish were raised to sexual maturity and kept under standard laboratory conditions at a temperature of 28°C and a 12:12h light-dark cycle. The fish were fed ad *libitum* three times per day with live *Artemia* (ZM Systems, UK) and dry food (Medium granular, ZM Systems, UK). The Swedish ethical standards were followed, and all experimental protocols were approved by the Swedish Board of Agriculture (Jordbruksverket, approval no. C 3/15).

### Experimental set-up for sperm sample collection

To collect sperm samples for sRNA-sequencing, males were exposed to one of two different treatments as described above and in (Zajitschek *et al*, 2014): (1) a high risk of sperm competition treatment (hereafter referred to as High) where two males were kept with one female, or (2) a low risk of sperm competition treatment (Low) where one male was kept with two females in 3L tanks. In zebrafish, both males and females compete for spawning and both sexes can be dominant (Spence *et al*, 2008). As larger fish tend to dominate smaller ones, we were careful to select fish with similar sizes for the experiment. To reduce stress exposure for experimental fish kept in such small groups, artificial aquaria plants were added to the tanks, providing sheltering and hiding space. The exposure duration corresponds to the time necessary for the completion of two spermatogenic cycles (Leal *et al*, 2009) and allows for any non-genetic signals to be potentially incorporated into newly produced mature sperm (Dias and Ressler, 2014; Rodgers *et al*, 2013).

Sperm samples were collected under anaesthesia (see Supplementary Information for details) from each male at two time points resulting in samples from High1 (High exposure first), Low2 (Low exposure second), Low1 (Low exposure first) and High2 (High exposure second) (Fig.1). Each treatment lasted for two weeks before sperm were collected. We conducted pairwise comparisons of Low treatments only (Low1.Low2), High treatment only (High2.High1) and between Low and High (Low2.High1, Low1.High1, High2.Low1, High2.Low2).

**Figure 1.**
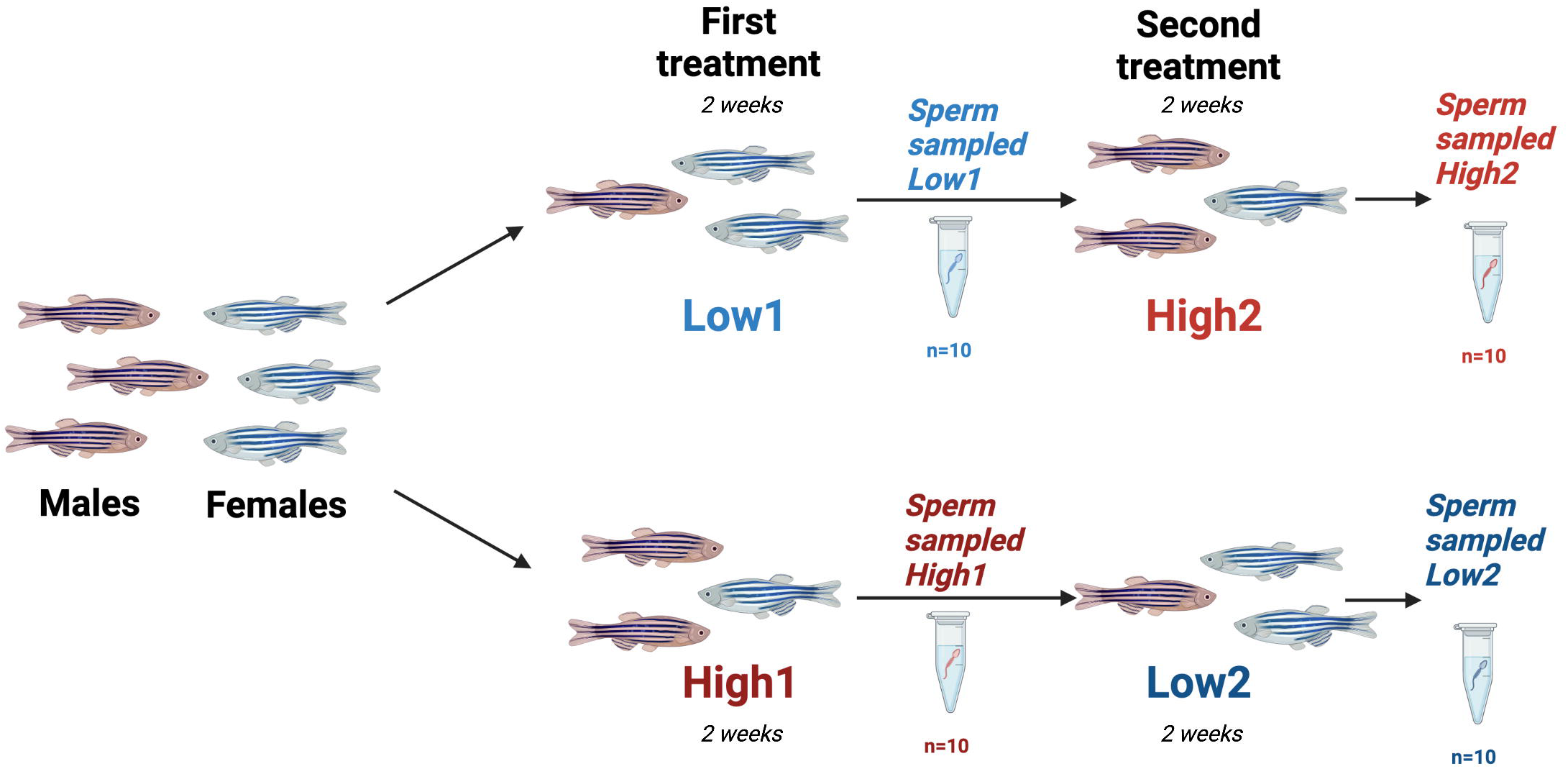
Schematic of the male competition environments in each treatment group for sperm sRNA sequencing experiment, namely a High competition treatment (two males and one female) and a Low competition treatment (one male and two females). All males were exposed to both treatments in randomised order with half starting High followed by Low and the other half *vice versa*. 1 indicates first treatment and 2 indicates second treatment. Sperm samples were collected after each treatment round for sequencing. We used n=10 males in each group resulting in a total of 40 sperm samples at the two time points. Red fish are male zebrafish, and blue fish are female zebrafish.

### Sperm phenotypic data analysis

Statistical analyses of the treatment effects on sperm traits were analysed using the package *lme4* (Bates *et al*, 2015) for the statistical software R, (see Supplementary Materials for details).

### RNA extraction, library preparation and sequencing for sperm sRNA analyses

RNA was extracted from individual sperm samples as described in *Supplementary Material*. Library preparation for sRNA from sperm samples was done using the New England BioLabs kit NEBNext^®^ Multiplex Small RNA Library Prep Set for Illumina^®^ Set 1 and 2 (NEB #E7300 and <u>NEB #E7580</u>). The provided guidelines of NEB were followed for small sample preparation until step 15, the performing of PCR Amplification. Eight individual sperm samples were sequenced per lane on Illumina HiSeq2500 with 50bp single-end kit resulting in an average read depth of 23 million reads per sample. Samples were combined to account for lane bias and excluding index-primer repeats. The raw sRNAseq datafiles are available on GEO here: GSE248535, https://www.ncbi.nlm.nih.gov/geo/query/acc.cgi?acc=GSE248535. Isolation of sRNA’s and quality checks can be seen in Suppl. Fig. S3 at the sequencing platform of SciLifeLab at Uppsala University.

### Experimental set-up for embryo sample collection

Experimental males were exposed to one of two different treatments as described above. After two weeks of exposure to treatments, the sperm of experimental males were collected and used for *in vitro* fertilisation (IVF) procedures as described in Supplementary Material. Females used for IVFs came from separate tanks and were not exposed to any experimental treatments but kept in a 1:1 sex ratio in 3L tanks and a density of 12 fish per tanks. These females will be referred to as neutral or control females. Embryos were collected at 24 hours post-fertilisation (hpf), a key stage in early embryo development with highly synchronised expression patterns (Mathavan *et al*, 2005), for whole transcriptome sequencing.

### Library preparation and sequencing for embryo transcriptome analyses

RNA extraction from embryos was performed as described in Supplementary Materials. Prior to library preparation, ribosomal RNAs were depleted by using the Ribosomal RNA Depletion kit (Ambion, A1083708) following the manufacturer’s protocol. cDNA libraries were prepared from RNA samples using the NEBNext Ultra Directional RNA Library Prep Kit (Illumina compatible, NEB, #E7420L following the kit protocol, generating 125 bp paired end reads. The RNA and cDNA libraries were prepared as described above from four individual embryos from each of the 20 sub-clutches and sequenced using Illumina HiSeq 2500 Systems. The raw RNA-seq data from embryos has been deposited at: https://www.ebi.ac.uk/ena/browser/view/PRJEB66218. There was an average read depth of 15 million reads per sample. The samples were sequenced at <u>the sequencing platform of SciLifeLab at Uppsala University</u>.

### Bioinformatic analyses

#### Sperm sRNA analyses

Small RNA reads were converted from FASTQ to FASTA format using *seqkit* v0.15.0 (Shen *et al*, 2016), and then processed to trim sequencing adaptors using a custom Perl script (Fowler *et al*, 2019)Supplementary Material) recognising the first eight bases of the adapter sequence (‘AGATCGGAAGAGC’).

For miRNA analsyes, the trimmed reads were aligned to miRBase (v22.0) mature zebrafish miRNA sequences using *PatMaN* v1.2.2, (Prufer *et al*, 2008), (parameters-e 0-g 0). A Perl script (Fowler *et al*, 2019) Supplementary Material) was used to parse the alignment files into a count table for each sample, which were then collated to generate an aligned read count table across all samples. *DESeq2* (Love *et al*, 2014) was used for normalisation of counts between samples and with treatment as the sole fixed effect to examine the effects of social stress. Adaptors for sequencing identified from FastQC reports and TrimGalore v0.6.5 were standard Illumina adapter: AGATCGGAAGAGC.

For piRNA analyses, the trimmed reads were aligned to the piRNA reference using *PatMaN* (Prufer *et al*, 2008), (parameters-e 0-g 0). A Perl script (Fowler *et al*, 2019) Supplementary Material) was used to parse the alignment files into a count table for each sample, which were collated to generate an aligned read count table across all samples. *DESeq2* (Love *et al*, 2014) was used for normalisation of counts between samples and calling differentially expressed piRNA clusters with treatment as the sole fixed effect to examine the effects of social stress.-Heatmaps of log^2^ fold change for differential expression of miRNAs and piRNAs were generated using the heatmap package version 1.0.12 in R Studio (Kolde, 2012).

#### Differential gene expression analyses in embryos

We sequenced a total of 80 embryos with four embryos per sub-clutch from a total of 20 sub-clutches from ten males (five per treatment) and ten females. The analysis of the RNA-sequencing data was conducted following (Conesa *et al*, 2016). Quality control was assessed using FastQC v0.11.6, (Andrews, 2010). Reads were mapped against the zebrafish reference genome version (Genome Reference Consortium, 2017) and counted at the gene level using STAR v2.5.3a (Dobin *et al*, 2013). Differential gene expression was analysed with the R package *DESeq2* v1.34.0, (Love *et al*, 2014) with treatment and female ID as fixed effect. We used R packages *EnhancedVolcano* v1.12.0 for plotting volcano plots (Blighe *et al*, 2021). ShinyGO v0.77 was used for GO terms analysis with a custom specified background list of all genes expressed with a *P* value in the *DESeq2* output results.

#### Sperm piRNA analysis

Zebrafish GRCz11 piRNA clusters were downloaded from piRNA Cluster Database (Rosenkranz, 2016) in bed format and GRCz11 reference assembly genome was downloaded from NCBI Genomes (https://www.ncbi.nlm.nih.gov/datasets/genomes/?txid=7955). A total of 532 piRNA cluster loci were extracted from the reference assembly genome using the genomic coordinates of the piRNA clusters in the.bed file using the *bedtools* (v2.29.2) function *getfasta*, producing the reference sequence for piRNA clusters. PiRNA loci not found in the NCBI reference assembly (e.g. loci in scaffolds/contigs) were not included.

#### De-novo sperm piRNA cluster generation

Small RNAs were adaptor trimmed using *trimgalore!* (v 0.6.6,--stringency 8),(Krueger, 2019), with cutadapt v1.18 (Martin, 2011). Trimmed sRNAs were then converted from fastq to fasta format using seqkit fq2fa (v0.15.0) (2) all sequencing files where concatenated into a single processed sRNAs fasta file for piRNA cluster prediction. To identify additional piRNA clusters not found in the piRNA cluster database, we ran the *proTRAC* pipeline v 2.4.4 on the fasta file of piRNAs generated as described above (Rosenkranz and Zischler, 2012). All sequencing files where concatenated into a single processed sRNAs fasta file for piRNA cluster prediction. In brief, the script TBr2_collapse and TBr2_duster was used to remove redundant and low-complexity sequences from the processed sRNAs. sRNAs were then mapped using the perl script *sRNAmapper* v1.0.5 (Roovers *et al*, 2015), to the *Danio rerio* genome GRCz11. The resulting map file was then processed with the Perl script proTRAC_2.4.4 to predict piRNA clusters. In total, we discovered 902 new piRNA clusters concatened with the existing 532 piRNAs from the reference genome, to generate a new piRNA reference with 1,434 piRNA clusters in total. Locus identifiers within the expression tables and results table starting with NC_are previously identifed clusters, all others are *de novo* identified with proTRAC. PiRNA loci located on unplaced scaffolds/contigs were not included.

The resulting fasta file containing piRNA clusters was concatenated with the existing piRNA database and used as the set of piRNA reference sequences called from this point onwards the ‘piRNA reference’ (1434 piRNA clusters in total). Locus identifiers within the expression tables and results table starting with NC_ are clusters from the existing database, all others are *de novo* identified with *proTRAC*.

#### Differential expression of embryo TEs

To identify differentially expressed TE families between the embryos of High and Low competition males, we used TEtranscripts v2.2.3 on the transcriptome data (Jin *et al*, 2015). The TETranscripts pipeline was run using-g GRCz11 genome settings. Strandedness of our RNA-seq data was checked with MultiQC report (Ewels *et al*, 2016). TETranscripts was executed with the following settings: *sorted_bam_files.bam--GTF Danio_rerio.GRCz11.107.gtf--TE GRCz11_Ensembl_rmsk_TE.gtf--mode multi--project projectname--minread 1-i 100--padj 0.05--outdir results*.

#### Sperm miRNA target gene analysis

For sensitivity, statistical confidence of miRNA-target gene associations in the 3’ untranslated region (UTR) of mRNAs by miRNAs, miRanda v3.3a pipeline was used with default settings (John *et al*, 2004). MiRanda was run with fasta file of mature zebrafish miRNAs and 3’ UTRs to scan and generate scored putative matches in the output. Full scripts and files can be accessed here: https://github.com/alicegodden/paternalsocstress/tree/main/miRanda. Zebrafish UTRs were obtained from TargetScan (Agarwal *et al*, 2015) and the mature miRNA fasta file was obtained from miRbase (Griffiths-Jones *et al*, 2006). The *ShinyGO* web-tool v0.77 was used for GO terms analyses on target genes (Ge *et al*, 2020). The background list of genes for sperm was taken from published scRNAseq datasets (Qian *et al*, 2022) and was limited to genes known to be expressed in sperm. To find and identify piRNAs in the differentially expressed clusters, the genomic regions were searched with piRBase (Wang *et al*, 2019).

To test for a link between sperm piRNAs and embryo TE, we used *FishPi*, a Python-based (*Python* v3.11) piRNA-TE complementarity analysis tool (Godden et al. currently under peer-review, can be found here: https://github.com/alicegodden/fishpi/). *FishPi* works by matching piRNA seed regions specified by user (1-10bp for 5’ end of mature piRNA for teleost) as input, and TE matches are identfied in a reference fasta file containing a list of all known TE sequences.

To link gene expression and TE expression in the embryo dataset, we developed Fish Transposable Element Analyser (*FishTEA).* We compared overlapping differentially expressed genes and TE loci using the DanRer11/GRCz11 reference genome. All scripts were written in Python v3.11. *FishTEA* works in five key steps that are ran individually: step 1: addition of TE loci co-ordinates, 2. Matching overlapping genes and TEs, 3. Generation of a chromosomal plot, 4-5 – plotting bar charts at TE family / class level count data. The aim was to visualise any overlaps between significantly differentially expressed genes and significantly differentially expressed TEs and to identify particularly active regions in the genome. *FishTEA* can be easily adapted to other organisms and can be found on Github: https://github.com/alicegodden/FishTEA/.

All scripts used in this project can be found here: https://github.com/alicegodden/paternalsocstress/.

## Results

### Phenotypic sperm traits

We found some significant differences in sperm traits between males from the High and the Low competition treatments, where Low competition males produced faster sperm than High competition males (Suppl. Fig. S1; Suppl. Table S1 for VCL).

### Differential expression of sperm miRNAs

Male ID had a small effect sperm miRNA profiles as highlighted by a Principal Component Analysis (PCA) across all samples where treatment was a secondary factor (Fig. 3A). We found a total of seven significantly differentially expressed miRNAs across the pairwise comparisons between the four treatment groups (Fig. 3B), one of which was found in the High2.Low1 comparison (*dre-miR-10b-5p*). This miRNA and the other 6: *dre-miR-129-5p*, *dre-miR-184*, *dre-miR-181a-5-3p*, *dre-miR-183-5p*, *dre-miR-193b-5p*, *dre-miR-10b-5p*, *dre-miR-200a-5p* were all significantly differentially expressed in the High2.Low2 pairwise comparison. No significantly differentially expressed miRNAs were found in any other pairwise comparison.

When looking at log_2_ transformed counts per million (CPM) Z-scores of significantly differentially expressed miRNAs across all samples (Fig.3C), miRNA expression showed minimal variance across samples, with Low2_6 and Low2_7 showing higher expression for all miRNAs. Low1 samples appeared to show the lowest expression of miRNAs, whereas High1 showing the highest expression of miRNAs.

### Differential expression of sperm piRNA clusters

We found a total of 1,434 piRNA clusters in sperm, of which 532 were already known (Fig. 4B), and 902 were unknown *de novo* clusters from the *proTRAC* output. A PCA of piRNAs showed some clustering of the individual treatment groups on PC1, with Low competition treatments clustering with some degree of separation from the High competition treatments (Fig. 4A).

We observed a general increase in expression of piRNA clusters in sperm samples of all Low males (Low 1 and Low2) compared to all High males (High1 and High2; Fig. 4B). However, we found significant differences when comparing the different treatments according to experimental order (High1 and High2 and Low1 and Low2). The most significantly differentially expressed piRNA clusters were found when comparing sperm samples from High2 and Low1 males, piRNAs in this comparison were mostly downregulated in expression. The most upregulated piRNA cluster in this comparison was *NC_007123.7:14899002-14913024* and the most downregulated clusters included many unannotatted piRNA clusters from *CHROM:14845000-14858022* to *CHROM:7007002-37018889*. A significant trend of downregulation of piRNA clusters was also observed in the comparison of groups Low1-High1, and High2-High1. The general pattern across these comparisons was the significant downregulation of piRNA clusters in sperm of males exposed to a High treatment (Fig. 4B).

Significanly differentially expressed piRNA read counts that were log_2_ transformed CPM with Z-scores showed higher piRNA expression in the High2 group, and lower overall expression in the High1 group (Fig. 4B). When looking for differential expression in piRNA clusters between treatments (Fig. 4C), we identified 45 differentially expressed piRNA clusters between samples from the same males in the comparison Low1.High2 (Fig. 4C, denoted by asterisks). Of note, there were no significantly differentially expressed piRNAs for the Low2.High1 and High2.Low2 comparison of samples. However, we found six differentially expressed piRNAs (Fig. 4C) in the Low1.High1 comparison, five differentially expressed piRNA clusters in the High1.High2, and one differentially expressed piRNA in the Low1.Low2 comparison (Fig.4C).

According to piRBase (Wang *et al*, 2019), piRNA cluster *CHROM: 57940016-57946585* is located on chromosome 1 and contains 125 known piRNA sequences within that cluster. This cluster was significantly downregulated in comparison Low1.High1. The largest identified piRNA cluster is located on chromosome 4, *NC_007115.7:52720448-52740603* at 20.16kb in size. It contains the sequences of 169 known piRNAs, and was significantly enriched in sperm from High competition males, showing the highest fold change in expression. The *NC_007133.7:32643430-32648896* cluster is located on chromosome 22, is 5.4kb in size and contains six known piRNA sequences, and three ncRNA genes: *BX640547.5*, *BX640547.4*, *BX640547.6*. This piRNA cluster was significantly downregulated in sperm from High competition males (High1 and High2), (Fig. 4B). From the largest piRNA cluster, *NC_007115.7:52720448-52740603*, individual piRNAs were further analysed with *FishPi* (Godden *et al*, 2023) to identify matching TE transcripts. As there were 169 piRNAs in this cluster, the first piRNAs of the three biggest subclusters were selected for analysis (Suppl Fig. 5). The most significant and complementary class of TEs to *piR-dre-43599*, *piR-dre*-*7058* and *piR-dre*-*58396* were class I DNA transposons, with fewer complementary class II RNA retrotransposons (Suppl. Fig. 5). *piR-dre-43599* was complementary to 2080 TEs (1846 Class I and 234 Class II – Suppl Fig. 5A), *piR-dre-7058* was complemenetary to 1160 TEs (874 Class I and 286 Class II-Suppl Fig. 5B) and *piR-dre-58396* was complementary to 2483 TEs (1943 Class I, 540 Class II-Suppl Fig. 5C).

### Differential gene expression in 24hpf embryos

We identified a total of 612 differentially expressed genes with a *p_adj_* < 0.05 between 24 hpf embryos sired by High and Low competition males (see Suppl. Files 3-4 for raw RNA-seq counts data and associated metadata). A PCA showed negligible clustering of expression profiles by treatment groups or maternal effects (Fig. 2A). Of the differentially expressed genes, 358 were downregulated and 254 were upregulated in High competition embryos. *ENSDARG00000104790* was the most downregulated differentially expressed gene in High competition embryos (Fig. 2C) and *Metazoa_SRP* (*ENSDARG00000103068*) was the most upregulated differentially expressed gene (Fig. 2C). GO terms analyses on biological processes showed strong associations of these two gene groups with muscle development and differentiation (Fig. 2B). *ENSDARG00000104790* (*CABZ01045617.*1) is expressed in the vascular system in zebrafish embryos (Gurung *et al*, 2022). *Metazoa_SRP* (ENSDARG00000103068), which is approximately 300 bp long (cDNA) and is present in multiple copies (81 hits on Ensembl) in the zebrafish genome, with most copies located on chromosome 11. These are necessary components for co-translational protein targeting by the signal recognition particle (Bradshaw and Walter, 2007; Keenan *et al*, 2001).

**Figure 2.**
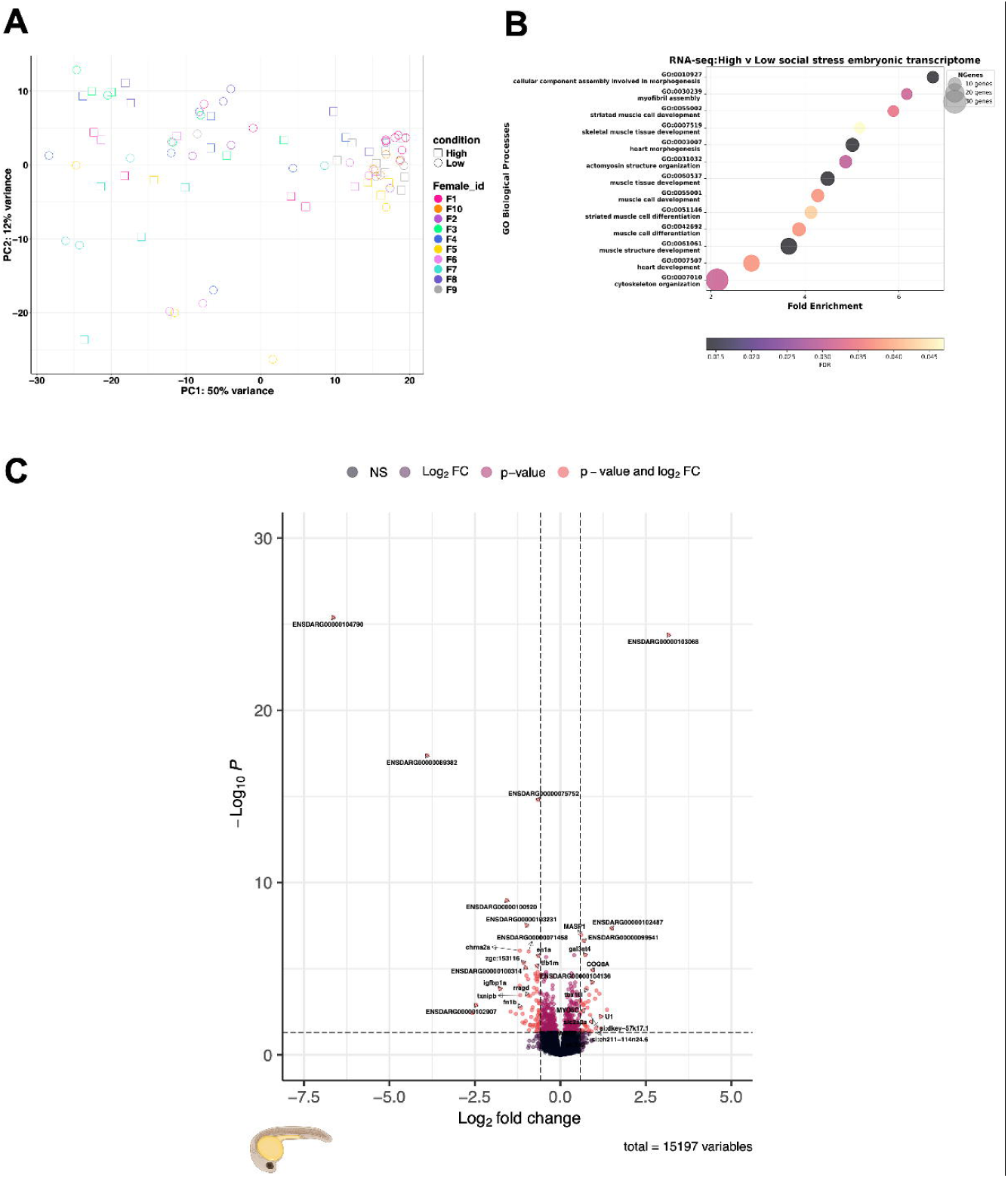
Differential expression of mRNAs in 24 hpf half-sibling embryos sired by males exposed to High and Low competition treatments. **(A)** Principal Component Analysis with treatment (Low v High), and female ID as factors from *DESeq2*. **(B)** GO term analysis looking at biological processes for zebrafish using *ShinyGO* on all significantly DE genes (FDR cut-off was set to *p_adj_* < 0.05) revealed a significant prediction to affect muscle development and differentiation. **(C)** Enhanced volcano plot of DE genes from *DESeq2*.

### Transposable element analyses

The embryo transcriptome data were analysed with the *TEtranscripts* pipeline (Jin *et al*, 2015) to profile differential expression of TEs at the family level (Fig. 5A). A PCA of the TE data showed modest clustering of the samples according to the two treatments and female ID (Suppl Fig. S4A). The most significantly differentially expressed TEs at family level were ERV, BEL and Gypsy TEs, which belong to the LTR class (Fig. 5A). The most abundant TEs were Class I DNA TEs (Suppl. Fig. S4B-C). We used *FishTEA* (Godden and Immler, 2024) to test if TEs overlapped with genes that were differentially expressed and to reveal regions of the genome showing enrichment or downregulation of gene expression or TE activity. We found 343 (out of 612) significantly differentially expressed genes to overlapp with 59 (out of 116) significantly differentially expressed TEs. We found that the majority of these TEs were Class I DNA TEs, as denoted by the orange dots, with a few Class II RNA retrotransposons, as denoted by the blue dots (Fig. 5B). Regions of highest enrichment for shared clustering of TEs and differentially expressed genes included the short arm of chromosome 4 and chromosome 23. Interestingly, the long arm of chromsome 4 showed little activity of significantly differentially expressed TE families and genes.

### Linking sperm sRNAs with DEGs in embryos

To link the significantly differentially expressed genes identifed in the embryos, and the significantly differentially expressed miRNAs in the sperm, we used *miRanda*, which identfies the genes targeted by specific miRNAs by searching for complementary hits in the 3’ UTR. The number of genes that are targeted by specific miRNAs are referred to as hits: *dre-miR-129-5p* had 140 hits, *dre-miR-184* had 30 hits, *dre-miR-181a-5-3p* had 22 hits, *dre-miR-183-5p* had 72 hits, *dre-miR-193b-5p* had 27 hits, *dre-miR-10b-5p* had 35 hits and *dre-miR-200a-5p* had 47 hits (Fig.3D). The 3’ UTR’s predicted to be the top hit of the significantly differentially expressed miRNAs showed 515 (out of 612) of the significantly differentially expressed genes as predicted targets (Suppl. Fig. S2H). We performed GO terms analysis to examine potential affected biological processes and found enrichment for terms associated with muscle development terms (Suppl. Fig. S6).

**Figure 3.**
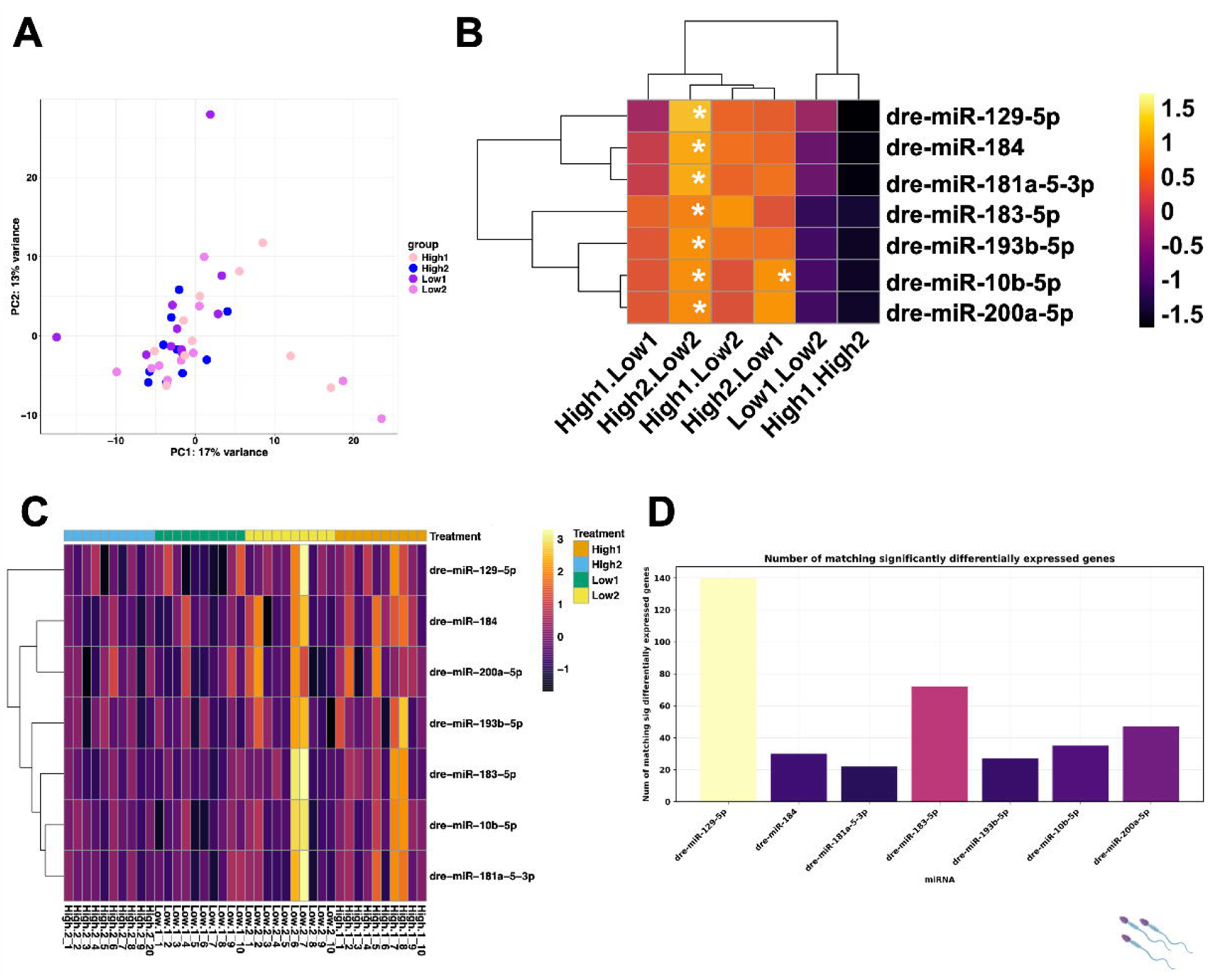
Differential expression of miRNAs in sperm of males exposed to High and Low male-male competition environments. **(A)** Principal Component Analysis showing variation across all samples used in the miRNA differential expression analysis. **(B)** Heatmap with dendrogram clustering showing all statistically significantly differentially expressed miRNAs log^2^ fold change (*p* = <0.05 denoted by *). Colour key reflects miRNA expression level, and shows general trend of miRNA expression. Not all miRNAs were significantly differentially expressed across all groups. **(C)** Heatmap of normalised counts (log^2^ CPM), from DESeq2 output of sRNA-seq data on miRNAs across all samples, grouped by treatment. **(D)** Number of matching significantly differentially expressed genes that targeted a significantly differentially expressed miRNA. 515 genes targeted the miRNAs.

We found 224 out of 612 significantly differentially expressed genes from our embryonic transcriptome data were targeted by the 7 significantly differentially expressed miRNAs from our sperm data, as predicted by miRanda (Suppl. Fig. S2A-G. In sperm from High2 males, the miRNA *dre-miR-129* was enriched (Fig. 3B) and was predicted to target significantly downregulated genes *ENSDARG00000075752* (*MYO18A*) and *ENSDARG00000103231* (novel gene) in the embryos (Suppl Fig. 2G). A Fisher’s exact test to examine for enrichment of miRNA target genes in the differentially expressed genes from embryos, showed that the differentially genes targeted by sperm miRNAs were highly enriched over random (odds ratio = 129.26; *P* = 2×10^-16^).

## Discussion

Variation in male-male competition not only affected sperm velocity parameters but also sperm sRNA, TE and gene expression profiles. Environmental changes and stress may affect the defense mechanisms in the male germ line where small RNAs play a key role in the defense of the genome against the activity of TEs and other possible DNA-damaging elements (Godden and Immler, 2023). These same signals may induce a potentially mild stress response in the resulting offspring, which affects early growth and development. Use of *IVF* in the externally fertilising zebrafish made our results powerful and allowed us to focus more on paternal effects while controlling for maternal effects (Mashoodh *et al*, 2018). Here, we discuss how a response to competitive male social environments may explain the changes in mi-and piRNA profiles in sperm and how these changes may be linked to peturbed gene and TE expression differences in the embryo RNA-seq data.

As previously described in many taxa (Bonduriansky and Crean, 2018; Zajitschek *et al*, 2017; Zajitschek *et al*, 2014), the level of male-male competition did have an effect on sperm velocity where low competition males produced faster sperm than High competition males (Suppl. Fig. S1; Suppl. Table S1 for VCL). This confirms that our treatments had the expected effects in response to male social environments, and that high competition males experienced more stress reducing overall sperm performance.

### Paternal stress effects on gene expression in zebrafish embryos

We found a significant general enrichment of GO terms for muscle development in High competition embryos (Fig. 2B). Specifically, T-box transcription factor 16-like gene *tbx16l* was enriched in embryos sired by males under High competition environments, a which is involved in mesoderm and somite development (Windner *et al*, 2015). An increase in expression of *tbx16l* could lead to more rapid muscle development and earlier onset of larval activity. Other significantly differentially expressed genes involved in embryo development include *si:dkey-121a11.3, myo18a* and *masp1*. Differentially expressed genes involved in the development of the central nervous system include *masp1*, *chrna2a* and *ern1,* and *en1a*. Finally, *gal3st4, ano7* and *coq8ab* are involved in general metabolic pathways and lipid metabolism. The GO terms for biological processes for miRNA revealed enrichment of processes in metabolism, muscle and nervous development (Suppl Fig. 5). All these processes indicate that embryos may differ in their developmental and metabolic rates, which links back to the earlier finding of increased hatching rates in embryos sired by High competion males (Zajitschek et al. 2014), and highlights potential detrimental efffects due to poor male condition in a high competition environment (Bonduriansky and Crean, 2018). Overall, we show that the response in males exposed to a High competition envrionments is translated into the offspring and triggers similar physiological responses. Such a response has been observed in offspring of starved zebrafish males where the gene expression in the embryos reflected gene expression changes observed in starved adult zebrafish (Jimenez-Gonzalez *et al*, 2024).

### Effects of paternal stress on TE activity in embryos

TE activity was altered at the family level, with strong enrichment and downregulation of TE families in High competition embryos (Fig. 5A). A possible explanation for the effects we observed is that males facing a rival male may have a physiological response to the higher stress levels induced by the competitive environment (Bonduriansky and Crean, 2018; Filby *et al*, 2010). This response affects the sRNAs in the sperm and their transfer into the zygote may explain the differential gene exrpession observed in the embryos as well as the differences in TE activity. Additionally, we found that 343 significantly differentially expressed genes overlapped with loci of 59 significantly differentially expressed TEs (Fig. 5B). These TEs may be hitchhiking the gene expression mechanisms to upregulate themselves, with two-thirds of zebrafish TEs known to be associated with host gene expression (Chang *et al*, 2022). Such detrimental TEs would be expected to be selected against and removed quickly over evolutionary times and time-point sampled transgenerational studies will give more insight into directional selection.

### Sperm miRNAs

SRNAs are known to be involved in inter-and trans-generational inheritance of parental conditions (Duempelmann *et al*, 2020; Rechavi and Lev, 2017). In zebrafish, at least 415 miRNAs are known, which are split into 44 famlies, with *miR-430* being the largest family with many isoforms; *miR-430* is located on chromosome 4, which has the least protein-coding genes of all zebrafish chromsomes (Howe *et al*, 2013; Thatcher *et al*, 2008). We found *miR-10b-5p* to be the most significantly differentially expressed miRNA in sperm of High versus Low competition males (Fig. 2b). This miRNA is highly conserved across the animal kingdom (Griffiths-Jones, 2004; Griffiths-Jones *et al*, 2006). In house mice, *miR-10b* is involved in spermatogonial stem cell production and loss of *miR-10b* may lead to increased spermatogonial stem cell death (Li *et al*, 2017). *MiR-10b* is targeting *Kruppel-like factor 4* (*Klf4*), a transcription factor that is involved in cell cycle regulation, contact inhibition and apoptosis, and is highly expressed in germ cells and Sertoli cells (Li *et al*, 2017). A possible explanation is that the high competition treatment led to changes in sperm production by deregulation of sperm stem cell renewal as shown in dominant zebrafish males that have higher numbers of spermatids in their testes in comparison to subordinate males (Filby *et al*, 2010). The trend of enrichment of miRNAs in response to environmental stress may represent an adaptive mechanism to fine-tune gene expression and preserve the germline genome (Godden and Immler, 2023).

Physiological stress affected sRNA profiles in sperm following heat-stress in *Drosophila* where genes coding for heat shock proteins, including *Hsp68* and *Hsp70*, became highly enriched, and some sRNAs (miRNAs and piRNAs), and Gypsy familyTEs were differentially expressed (Bodelon et al, 2023). Similarly, *Drosophila* sperm miRNA profiles changed in response to variation in population sex ratio, where *miR-184* was found to be differentially expressed (Hotzy *et al*, 2021). In fruitflies, *miR-184* is important in the production of the female germline (Iovino *et al*, 2009) and given that mature *miR-184* is conserved between zebrafish and fruitflies, it may have shared functions in germ cell development (Zhang *et al*, 2023). Other differentially expressed miRNAs in our study included *dre-miR-129-5p, dre-miR-181A-5-3p and dre-miR-193B-5p,* (all enriched in High2.Low2). *Dre-miR-129* is conserved across vertebrates and is involved in zebrafish ciliogenesis, with inhibition of *dre-miR-129-3p* showing defective embryo development and suppression of ciliation in the Kupffer’s vesicle (Cao *et al*, 2012). *Dre-miR-181A-5-3p* is involved in zebrafish vascular development where experimental over-expression and knock-down impairs blood vessel development (Ma *et al*, 2019). Additionally, *miR-181A-5p* is known to regulate inflammatory immune responses and is involved in tumour development (Ye *et al*, 2018). Finally, *dre-miR-193B-5p* was found to be significantly downregulated in zebrafish larvae exposed to 1% ethanol (Soares *et al*, 2012). Stress induced by the high competition environment may therefore affect immune response and cardiac and developmental pathways. *Dre-miR-200* was enriched in High versus Low comparison groups, most significantly in High2.Low2. The *dre-miR-200* cluster is involved in the regulation of sperm motility, with increased expression leading to decreased sperm motility by targeting sperm-motility genes including *amh* (Xiong *et al*, 2018).

To link sperm miRNA with the embryo transcriptomes, the significantly differentially expressed miRNAs were put through miRanda to analyse predicted gene targets (Suppl. Fig. S2). Of the 612 significantly differentially expressed genes from our embryonic RNA-seq data, 515 were targeted by our miRNAs, with each miRNA targeting some significantly differentially expressed genes, enrichment testing with Fisher’s exact test showed *P* = 2.2 x10^-16^ with an odds ratio of 129.256. This shows a strong enrichment of the miRNA target genes (Fig. 3D). Each of these miRNAs were enriched following their competition environment treatment, in the High2 v Low2, High2 v Low1 comparisons (Fig.3B).

### Linking sperm piRNAs and embryo TEs

PiRNA enrichment helps to maintain the integrity of the germline through complementary binding to transposon transcripts, whereas loss of piRNAs leaves the germline genome vulnerable to invasion and disruption by transposons (Aravin *et al*, 2007). Chromosome 4 is the putative sex chromsome in zebrafish and hosts genes that can influence sex determination, particularly around position 61,176,889 (Anderson *et al*, 2012; Howe *et al*, 2013), which may explain the over-representation of repetitive elements in that region. We found the largest DE piRNA cluster *NC_007115.7:52720448-52740603* on chromosome 4 to be enriched in High competition sperm (Fig. 4B-C), which supports the idea that piRNA expression is linked to a response in the germ line to environmental stress to protect the germline genome (Godden and Immler, 2023; Klattenhoff and Theurkauf, 2008). No significantly differentially expressed genes and overlapping TEs on a significant portion of the long arm of chromsome 4; a region that is known to be particularly rich in piRNAs and in retroelements (Chang *et al*, 2022; Howe *et al*, 2013).

**Figure 4.**
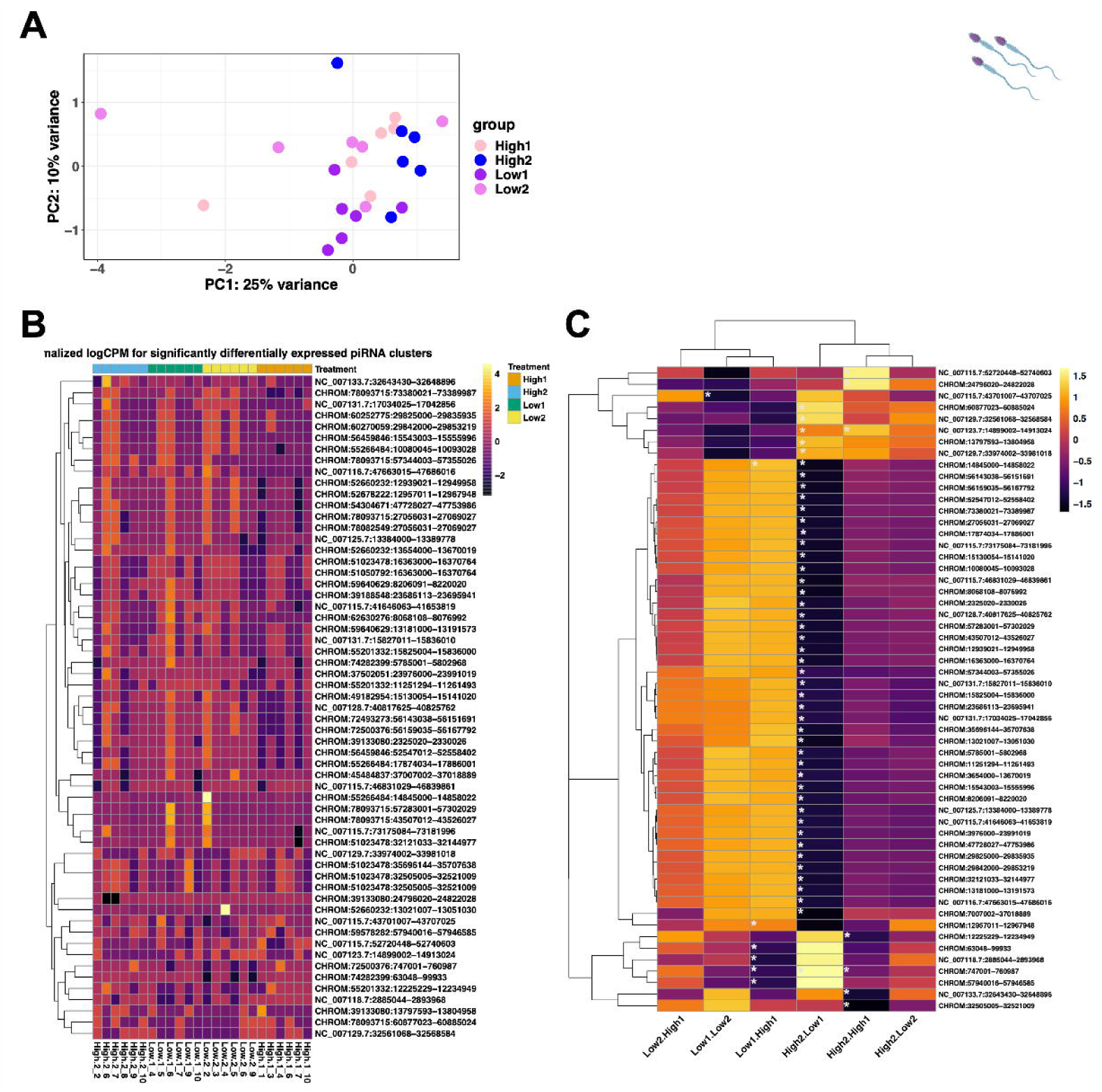
Differential expression of piRNAs in sperm of males exposed to High and Low male-male competition environments. **(A)** Principal Component Analysis showing variation across all samples used in the piRNA differential expression analysis. **(B)** Heatmap of normalised counts (log^2^ CPM) from DESeq2 output of sRNA-seq data on piRNAs across all samples, grouped by treatment. **(C)** Heatmap of all significantly DE piRNAs clusters log^2^ fold change (*p* = <0.05 are denoted by *) for pair-wise comparisons. Colour key shows piRNA expression level, and shows general trend of piRNA expression. PiRNA names are abbreviated (see Suppl. File S1 for full piRNA cluster loci names).

Approximately 56-59% of the zebrafish genome consists of TEs (Shao *et al*, 2019), and the majority are DNA transposons covering 46% and RNA retrotrotransposons 13% respectively (Chang *et al*, 2022). We found LTR elements to be most significantly differentially expressed between High and Low embryos (Fig. 5A). LTRs are most enriched during early embryonic developmental stages (Chang *et al*, 2022), which may explain a bias towards the significantly differentially expressed LTR elements in our study. Most zebrafish piRNAs map to TEs (Houwing *et al*, 2007), where a majority map to LTR elements, and fewer piRNAs map to DNA transposon targets, suggesting that piRNA changes with the environment to maintain the integrity of the germline genome (Chang *et al*, 2022). Our results suggested substantial complementarity of some of piRNAs in the differentially expressed cluster *NC_007115.7:52720448-52740603* to DNA TEs (Suppl Fig. 6). DNA TEs are linked to older TE age and events, (Chang *et al*, 2022) and retained activity correlated with longevity in *C*. *elegans* (Sturm *et al*, 2023). Therefore it is expected that piRNAs complementary to the LTR elements are responding to increased TE activity from the environmental stressor.

**Figure 5.**
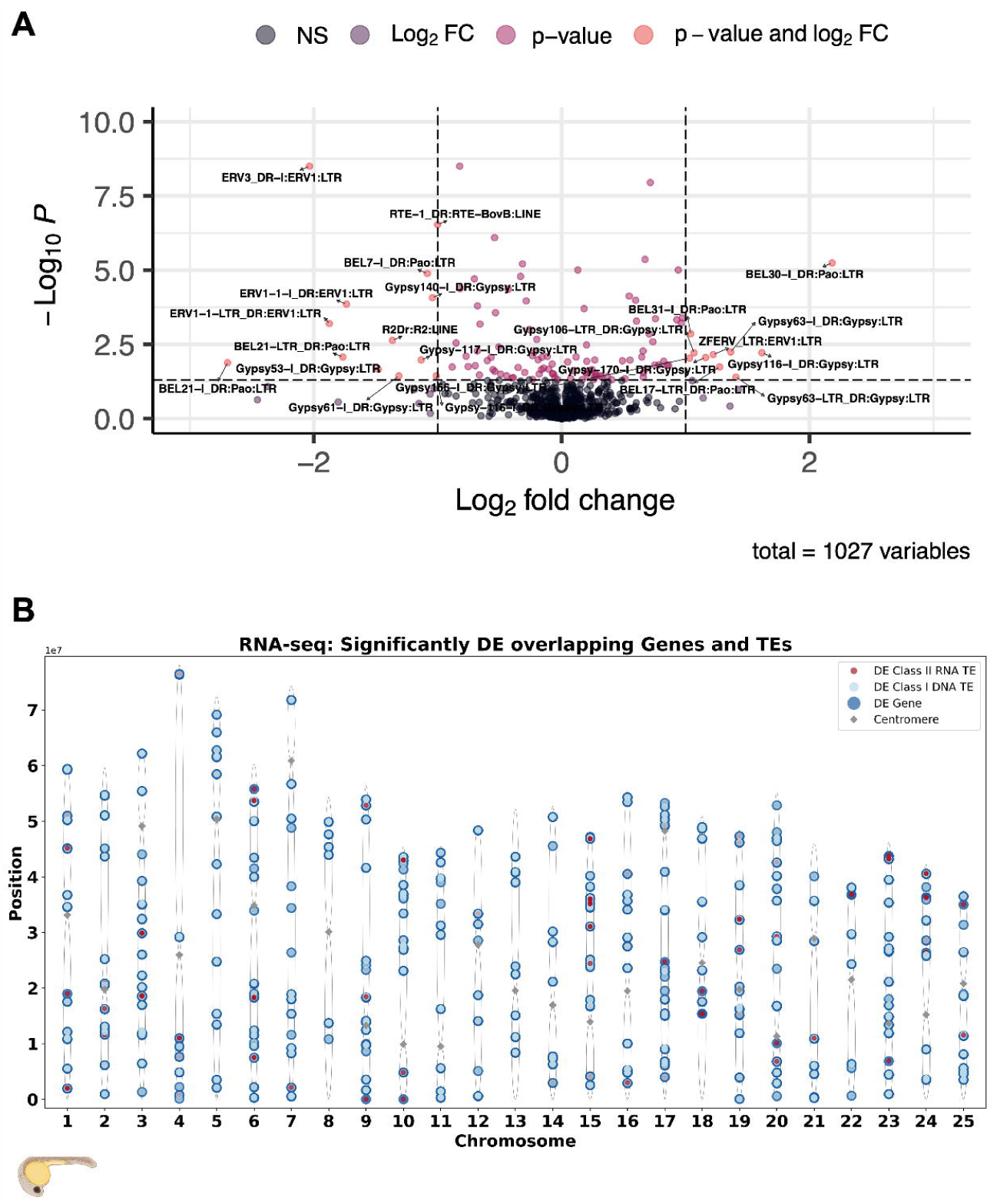
Differential expression of TE transcripts by class and family in embryos fathered by males from High versus Low competition environments. **(A)** Volcano plot of differentially expressed TEs by family from *DESeq2* analyses looking at impact of high social stress and factoring in female_id. **(B)** Chromosomal plot displaying the significantly differentially expressed genes and TEs that overlap along each chromosome: DE Genes (dark blue), DE TEs Class I (red), DE TEs Class II (light blue).

In the zebrafish genome, LTRs and LINEs are mainly located in pericentromeric regions whereas DNA transposons are located near or within genes; with over 60% of TE transcripts thought to be driven by local gene promoters (Chang *et al*, 2022). In the adult zebrafish genome, 37% of TEs are located in active regulatory states (Lee *et al*, 2022). In accordance, we found significantly differentially expressed TEs from Class II retroelements in the pericentromeric region, and Class I DNA TEs overlapped with gene loci more abundantly and frequently than Class II elements (Fig. 5B, Suppl Fig. 6). This finding suggests that DNA TEs may be expressed more readily, and co-expressed with genes expression resoonding to the High competition environment.

Many piRNA functions remain unknown, and most but not all piRNAs currently match transposon sequences. In addition, multiple piRNAs may target the same or different TE mRNAs (Zhang *et al*, 2018). Therefore, continued development of bioinformatic tools such as FishPi (Godden *et al*, 2023), FishTEA (in this paper), and CRISPR of TEs (Guo *et al*, 2024) as well as functional tests by knocking out mi-and piRNAs are needed to further analyse the function and interactions of sRNAs and TEs to further understand the mechanisms of their response to envionmental triggers.

### Social dominance and environment change

When running pairwise comparisons of sperm sRNAs between specific treatment groups (Low1, Low2, High1, High2), we identified a number of differentially exressed mi-and piRNAs that varied across comparisons When comparing Low1 and Low2 ony, we observed general downregulation of miRNAs and enrichment of piRNAs, whereas when comparing High2 and High1, we observed downregulation of miRNAs and piRNAs. In sperm from males in a Low treatment followed by High exposure, we found modest enrichment of miRNAs and some enrichment and downregulation of piRNAs. In sperm of males exposed to High competition, we found modest enrichment of miRNAs and downregulation of piRNAs. Overall, exposure to the High competition treatment seemed to have the most profound effect and led to significant changes in sRNA expression, and males exposed to High competition first and then Low competition differed from males, that had only been exposed to a Low competition ennironment. Nevertheless, the environment a male was exposed to immediately before sperm collection had the strongest effect on the direction of mi-and piRNA expression. It appears therefore that the history and specific sequence of environments affects several spermatogenic cycles. Spermiogenesis takes approximately one week in zebrafish (Leal *et al*, 2009), so having had two cycles of two weeks the second sperm sample is still showing some signs of impact from the first environmental exposure.

Thermal stress environments have previously been shown to reduce male fertilisation success in zebrafish, with increasing thermal stress negatively affecting sperm quality and subsequently fertility (Irish *et al*, 2024). Similarly, alarm cues affect sperm sRNAs in zebrafishsupporting the idea that the paternal environment may influence offspring phenotype, via paternal inheritance of sRNAs (Ord *et al*, 2020). Our study showed that 224 of 612 significantly differentially expressed genes were predicted targets of the significantly differentially expressed miRNAs. Together, this highlights the importance of the role of miRNAs for paternal inheritance.

## Conclusion

We provide clear evidence that variation in the social environment of zebrafish males can affect sRNA expression in their sperm and gene TE expression in the resulting offspring. We provide direct links between sperm sRNAs and differentially expressed gemes in the embryos by introducing new bioinformatic tools and link the seven significantly differentially expressed sperm miRNAs to a global regulatory function over the 612 signficantly differentially expressed genes in the embryos. Overall, we provide links between genes and TEs (FishTEA), genes and miRNAs, piRNAs and TEs (FishPi). The regulatory functions of many of the differentially expressed genes are in line with the general phenotypic effects described in previous studies (Zajitschek *et al*, 2017; Zajitschek *et al*, 2014). We are still at the very beginning of understanding the potential effects of paternal condition on offspring fitness, but our study further supports the idea that variation in paternal condition, even over short periods of time, is likely to have a significant effect. We currently can only speculate about the mechanisms responsible for the transfer of signals about paternal condition from father to offspring (Immler, 2018; Krawetz, 2005) and understanding these mechanisms certainly deserves more attention in the future. Whether the epigenetic changes caused by a environmental variation are a true inheritance factor or rather a marker of underlying mechanisms defending the germ line genome (or both) remains to be further investigated.

## Author contributions

AMG – Bioinformatic analysis, FishTEA creation and development, sRNA and mRNA data analysis, manuscript writing and preparation. WTAF – experimental design, running of experiments, RNA-sequencing analyses, manuscript writing. BK – experimental design, running of experiments, sRNA sample collection and preparation. LF – Bioinfomatic analysis of sRNA data. GA & CJ – Experimental work. SI – Planning of study, experimental design, funding acquisition and manuscript writing.

## Abbreviations

CPM: counts per million
FishTEA: fish transposable element analyser
hpf: hours post fertilisation
IVF: in vitro fertilisation
LTR: long-terminal repeat
miRNA: microRNA
piRNA: piwi-interacting RNA
sRNA: small non-coding RNA
TE: Transposable element
UTR: Untranslated region
VAP: average path velocity
VCL: curvilinear velocity
VSL: straight line velocity

## Acknowledgments & Funding

We thank Oded Rechavi and Sarit Anava for advice on the sRNA extractions and to Oded Rechavi for valuable comments on an earlier draft of the manuscript. WTAFS was funded by a Sven and Lilly Lawsky PhD scholarship and the study was supported by grants to SI from the Swedish Research Council, the European Research Council (ERC, Hapsel336633), the Human Frontier Science Program (HFSP R0025/2015), the Knut and Alice Wallenberg Foundation, the Natural Environment Research Council (NE/S011188/1) and the European Research Council (SELECTHAPLOID - 101001341).

## Data archiving

The raw sRNAseq datafiles are available on GEO: GSE248535, https://www.ncbi.nlm.nih.gov/geo/query/acc.cgi?acc=GSE248535 (under embargo until publication, can provide reviewer link upon request). The raw RNA-seq datafiles are available on the European Nucleotide Archive (ENA, https://www.ebi.ac.uk/ena/browser/view/PRJEB66218). All metadata, raw counts data, and scripts used to generate and visualise the results used in this project are available as supplementary files in the supplementary materials.

## Research ethics statement

All experiments described here were performed in accordance with the guidelines and approved by the Swedish Board of Agriculture (Jordbruksverket approval number C341/11).

## Supplementary Files

**Supplementary materials, methods and figures and figure legends**

Supplementary Material_heredity.docx

**Supplementary File S1- List of piRNA cluster full names in Fig. 4B**

Supp_File_1_piRNAs_heatmap_fullnames.xlsx

**Supplementary File S2 – Phenotypic data**

Supp_File_6_sperm_motility_data.csv

**Supplementary File S3 & S4- raw counts and metadata file RNA seq**

Supp_File_3_RNA_seq_rawcounts.csv

Supp_File_4_rnaseq_metadata.csv

**Supplementary File S5 & S6- raw counts and metadata file sRNA-seq: miRNA data**

Supp_File_5_40_mirna_counts_LF.xlsx

Supp_File_6_40_mirna_metadata_LF.csv

**Supplementary File S7 & S8- raw counts and metadata file sRNA-seq: piRNA data**

Suppl_File_7_piRNA_raw_counts.csv

Supp_File_8_pirna_metadata.csv

**Supplementary File S9 & S10- TEtranscripts RNA-seq raw counts output and metadata file**

Suppl_File_9_tetranscripts_raw_counts_Teonly.xlsx

Suppl_File_10_metadata_rnaseq_TEtranscripts.csv

