## Supplementary Materials Methods Figures for "Environmentally induced variation in sperm sRNAs is linked to gene expression and transposable elements in zebrafish offspring"

**Supplementary Material**

**Material & Methods**

*Statistical analyses of sperm phenotypic data*

We investigated the treatment effect on the curvature of three sperm velocity traits, namely VSL (straight line velocity), VCL (curvilinear velocity), and VAP (average path velocity), with social treatment, status, and time (in seconds post-activation, spa) as fixed effects, and male identity as a random effect on all time terms. We modeled the effect of time as a third-order orthogonal polynomial and included all interactions between treatment, status and time into a full model and removed them stepwise if non-significant. The analyses were performed for each sperm trait individually on a dataset containing 86 experimental males, including the ten males used in the fertilisation experiment (embryo transcriptomes). Additionally, after stepwise removal of non-significant interaction terms, the resulting models for each sperm trait were applied to the dataset containing only the ten males used in the fertilisation experiment. *P*-values for sperm trait curves were obtained by normal approximation (assuming the *t*-distribution converges to the *z*-distribution). Differences in fertilisation success (number of fertilised eggs bound to number of unfertilised eggs per half-clutch) between social treatments were analysed using a binomial generalised linear mixed model with social treatment as a fixed effect and male and female identities as random effects.

*Sperm sRNA extraction and library preparation for sequencing*

For sRNA extraction, initially total RNA was extracted and then sRNAs were separated later via a gel-run. Visibility of the material is enhanced via usage of Glycogen. Note that all solutions should be prepared with nuclease-free water, as no RNase inhibitor is used in the protocol. Prior to use, all equipment was thoroughly cleaned with RNase-away (Thermo Scientific™ RNase AWAY™ Surface Decontaminant Spray). Gloves (double layer) are mandatory throughout the entire extraction process to avoid any cross contamination and ensure personal safety according to chemical safety regulations. Because no digestive step was added, cleanliness and an RNase-free working environment including RNase-free reagents are essential for the success of the extractions.

Samples were removed from their storage at -80°C and left to thaw on ice. 400ul of TRIzol® Reagent was added followed by careful homogenization of the samples at room temperature using a micropestle and a handheld pestle motor until complete dissolution. Each sample was then treated with three different sizes of needles attached to a syringe (first 25G, then 27G and finally 30G). The entire sample was loaded and ejected from the syringe five times with each needle size to break the sperm membrane, loosen the nucleus and release the trapped RNA molecules. After the syringe step, samples were left standing for 3 minutes at room temperature.

*Phase separation, phase lock and washing for sperm sRNA*

400 µL of chloroform was added, the tube capped securely and vigorously shaken for 15 seconds after which it was left to incubate for 3 minutes at room temperature. This step was followed centrifugation at 16,000 rcf for 5 minutes at 4°C, which resulted in the separation of three phases: a lower red phenol-chloroform phase, an interphase and a colour-less upper aqueous phase containing the RNA. The aqueous phase should form about 50% of the total volume. The upper aqueous phase including the interphase was carefully removed using a pipette by twisting the tube at a 45° angle and transferring the supernatant into a pre-spun Heavy phase lock tube (2 mL tubes 5Prime Heavy Phase lock gel, Quantabio). In a next step, 1 mL Chloroform per 1 mL aqueous phase was added. This procedure was performed under a ventilation hood at room temperature. The resulting mix was inverted ca. five times until an emulsion forms, which is immediately centrifuged at 16,000 rcf for 5 minutes at room temperature. The resulting aqueous layer was transferred to a clean Axygen tube as used above.

*RNA precipitation on sperm sRNA*

Measuring the aqueous phase, the equivalent of 1/10 of the volume of NaOAC (3M, pH = 5.2) and the equivalent volume of 1/1 of isopropanol was added to the sample tube and transferred to the -80°C freezer overnight. The next day, the samples were thawed on ice and centrifuged at 16000rcf for 30 minutes at 4°C. The supernatant was carefully removed with a pipette leaving the white pellet at the bottom of the tube. The pellet was washed with 200µl cold (from -20°C freezer) 70% EtOH and mixed by gently inverting the tube to avoid moving the pellet and then left for 20 minutes standing at room temperature. After that, the sample was centrifuged at 16000rcf for 10 minutes at 4°C. All EtOH was then carefully removed, and the washing step repeated as above, but only left to stand for 5 minutes at room temperature. After centrifuging at 16000rcf and 4°C all EtOH was carefully removed (to remove all droplets a quick spin with a Microcentrifuge, MiniStar silverline, VWR was included) and the pellet air-dried for 5-10 minutes at room temperature (monitor drying process). The resulting pellet was resuspended in 8ul of pre-warmed to 70°C DEPC treated water, 2µl of the suspension were directly transferred into a TapeStation tube (Agilent Optical tube strip) for quality control, the remaining 6 µL were stored for a maximum of 1 week at -20°C for Library preparation.

*Quality control*

A quality control of the extracted RNA was performed utilizing the above prepared TapeStation tubes and Agilent High Sensitivity RNA ScreenTape and reagents (Suppl. Fig. 3)

*Preparation of sperm RNA sequencing pools*

Eight individual sperm samples were combined into pools for one lane each for sequencing on Illumina HiSeq2500 with 50bp single-end kit resulting in an average read depth of 23mio reads per sample. Samples were combined to account for lane bias and excluding index-primer repeats. Using 40µl of the sample with the lowest pM/µl, the samples’ total molarity in the pool was calculated (40* x pM/ul). The amount added from the seven remaining samples of the pool was calculated to fit this molarity (molarity sample 1/molarity sample 2 = amount sample 2 in pool). The assembled pools of total RNA were run on a 4% low melt agarose gel to separate target bands for size selection of targeted small RNAs.

*Sperm sRNA size selection in gel*

Small RNA was separated from total RNA using the E-Gel EX iBase system from Invitrogen. Cassettes of 4% low melt Agarose Gel were used with an E-GEL 50BP DNA ladder for this gel (Invitrogen). The gel wells were filled with 20µl ladder on the two outer lanes and 20-25µl of the sample pool in the central wells. The amount added per well depended on the volume of the pool, non-used wells were filled with ladder/ water for optimal run. The cassette was opened, and target bands cut out and transferred into a fresh and weighted 1.7ml Axygen maxiclear tube as described above. The weight of the cut bands with gel was recorded. To separate the target bands from the gel we used the Qiagen MinElute Gel Extraction kit.

*Sperm sRNA Target band separation from Agarose gel*

To the Axygen tube containing the gel, 3 gel-volumes of QG buffer is added. The mixture is incubated at 50°C for 10 mins, to facilitate full dissolving of the gel-band-mix, the tube is inverted every 2 minutes. It should turn into yellow fluid when fully dissolved. At full fluidity 1 gel volume of Isopropanol is added and invert mixed. In case of too large gel volume, it may be split into 2 tubes to continue. The sample is transferred into MinElute Spin columns 750 µL at a time and centrifuged for 1min at 13,000rpm and room temperature (RT). Flow-through is discarded and the filtering repeated in the same column until the entire sample from one gel is filtered. Following this 500ul QG Buffer is added to the MinElute Spin column and centrifuged for 1min at 13, 000rpm at RT. Flow-through is discarded and the collection tube can be re-used. 750 µL PE Buffer is added to the MinElute Spin column and left to incubate at RT for 5mins. The spin column is centrifuged for 1min at 13 000rpm RT, flow-through discarded and collection tube re-used for the next step. The “empty” collection tube is centrifuged with closed lid for 1min at 13,000rpm RT to ensure filtering out of the entire volume of PE Buffer. Flow-through and collection tube are discarded. The spin filter is placed in a fresh 1.7 mL low retention tube (Axygen Maxyclear Boil-Proof, Maxymum Recovery® tube) and 12 µL ultra-pure ddH_2_O. The filter with the added water will be incubation for 1min at RT, followed by centrifugation for 1min at 13,000rpm RT. The elute can be stored in -20 °C until sequencing preparation/ delivering to a sequencing platform.

*Sperm collection and velocity measurements for embryonic transcriptome project*

Twenty-four hours before the IVFs, the experimental males where physically separated from their company females by transferring the females into a breeding unit, which was kept inside the original tank. Males could therefore still receive the visual and olfactory cues from the females but could no longer interact. For the collection of the ejaculates, experimental males were anesthetized in MS-222 (Sigma-Aldrich, A5040) solution (0.016%) for a maximum of two minutes, rinsed in system water to remove excess MS-222 and placed ventral side up into a moist sponge under a stereomicroscope (Nikon SMZ800 C-DS). The genital pore was carefully blotted dry with a paper towel prior to sperm collection to avoid premature sperm activation upon contact with water. For each individual, sperm were collected using a calibrated micro-capillary (Sigma-Aldrich, P0674), transferred to an Eppendorf tube containing with 30μL of Hank’s buffer (recipe described by (1)) and placed on ice until measurement of density and velocity parameters (straight line velocity, VSL; curvilinear velocity, VCL; average path velocity, VAP). Sperm density and velocity were recorded using a brightfield microscope (UOP UB200i, PROISER R+D) equipped with a heating plate at 28°C (Linkam DC95, Surrey, UK) and a camera (ISAS 782M, PROISER R+D). The ejaculates were gently homogenized and 0.5 μL of the sperm solution was placed on a 20μm glass cell and activated with 4μL of system water at 28°C. The sample was covered with a coverslip and the density and motility recorded every 10 seconds for one minute, starting 10 seconds after activation. The measurements were repeated at least three times for each sample. The qualitatively suitable recordings were analysed using computer assisted software analysis (ISAS Proiser, Projectes I Serveis R+D, S.L., Valencia, Spain) with the following settings: frame rate of 50 frames/second, 50 frames used, particle area of 5-50μm^2^, threshold measurements for VCL values (μms^-1^): static<10<slow<45<medium<100<rapid.

*Egg collection for embryonic transcriptome project*

Females used for IVFs came from separate tanks and were not exposed to any experimental treatments but kept in a 1:1 sex ratio in 3L tanks and a density of 12 fish per tanks. These females will be referred to as neutral or control females. For egg collection, females were anesthetized in MS-222 for a maximum of two minutes, rinsed in system water and gently dried on a clean paper tissue. Eggs were collected into a clean, dry Petri dish by gently squeezing the female’s belly towards the genital pore. Contact of the eggs with water was carefully avoided to prevent premature activation.

*In vitro fertilisation embryonic transcriptome project*

Clutches with good quality eggs (i.e. yellow and translucent eggs) were divided into two sub-clutches and placed at opposite sides of the Petri dish. Each sub-clutch was fertilized with 10μL of sperm-Hank’s buffer mix by one male and resulting in one sub-clutch being fertilized by a high treatment male and the other by a low treatment male. Each experimental male fertilized sub-clutches from two neutral females (Figure S1). Such a split-clutch design allowed us to account for putative female effects. A total of 20 half-clutches were obtained from 10 males (five per treatment) and 10 females.

The developing embryos were kept in 150mm x 15mm Petri dishes containing methylene blue diluted in system water and incubated at 28°C. The fertilization success (number of fertilized eggs vs. total number of eggs) and survival at 2 hours post-fertilization (hpf) were recorded. Dead or abnormal embryos were recorded and removed. Four 24 hpf individual embryos from each half-clutch were frozen in liquid nitrogen and stored at -80°C until RNA extraction.

*Embryo RNA extraction for embryonic transcriptome project*

Total RNA from four frozen individual embryos at 24 hpf per sub-clutch was extracted. In total, RNA was extracted from 80 individual embryos. Prior to extraction, 100 μL of freshly made lysis buffer (1M Tris buffer, pH=7.5, 0.5M EDTA, 1M NaCl, 10% SDS, 20 mg/mL Proteinase K, was added to each sample. The tissues were disrupted mechanically by pipetting up and down and incubated in lysis buffer for 10 minutes. During incubation, the tubes were inverted gently every two minutes to ensure homogeneity of the samples. The samples were treated with 2 μL of DNase I (Invitrogen, 18068015) for 15 minutes at room temperature to remove any traces of genomic DNA. Total RNA was then extracted by adding 100 μL of phenol:chloroform:isoamyl (pH=4.5; Sigma-Aldrich, 77617) and 0.5 μL of glycogen (20mg/mL stock, 20 µg final Roche, 10901393001) to each tube. The samples were homogenized by gently inverting the tubes and centrifuged at 16,000 rpm for 15 min at 4°C. The upper aqueous phase was collected and transferred to phase-lock tubes, and mixed gently with 100 μL of chloroform:isoamyl alcohol (Sigma-Aldrich, C0549) to remove traces of phenol and ensure the purity of the RNA. The phase-lock tubes were centrifuged at 16000 rpm for 15 minutes at RT and the upper phase transferred to a new clean Eppendorf tube.

For RNA precipitation, samples were incubated for 30 minutes at -80°C with 1/10 volume of NaOAC (3M, pH=5.2, Thermofisher, R1181) and 1/1 volume of isopropanol (Sigma-Aldrich, I9516). After incubation the samples were centrifuged, and the upper phase carefully removed.

*RNA quality control*

The integrity of each sample was verified using the Agilent TapeStation system and High sensitivity RNA ScreenTape kit (Agilent Technology, 5067-5579). The samples were prepared following the kit instructions. Briefly, 2 μL of RNA sample or RNA ladder were mixed with 1 μL of High Sensitivity RNA sample buffer and vortexed for one minute. The samples were then incubated at 72°C for 3 minutes and transferred to ice immediately. RNA samples were loaded into the 2200 TapeStation instrument. The TapeStation generated an RNA integrity number (RIN value) for each sample (Agilent Software packages). Only samples with a RIN value higher than 8 were used to generate cDNA libraries.

**Supplementary Figures**

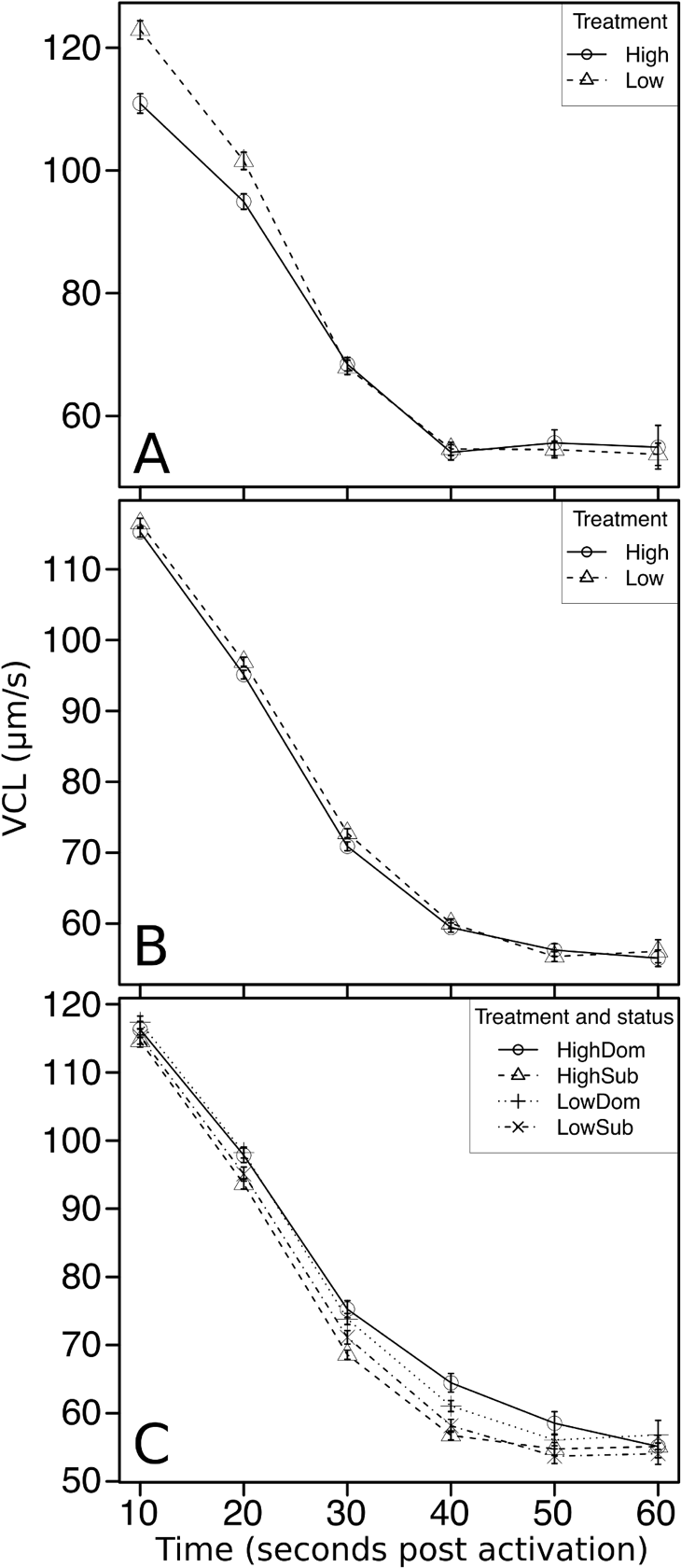

**Supplementary Figure 1.** Change in sperm VCL over time in seconds after sperm activation. (**A**) Sperm VCL per social treatment for 10 males used in embryo transcriptomics analysis. (**B**) Sperm VCL per social treatment for 86 males, including the 10 of the males used in embryo transcriptomics analysis. (**C**) Sperm VCL per social treatment and status for 86 males, including the 10 males used in embryo transcriptomics analysis. HighDom is for high treatment and status as dominant, HighSub is for high treatment and status as subordinate, LowDom is for low treatment and status as dominant, LowSub is for low treatment and status as subordinate. Whiskers indicate 95% confidence intervals.

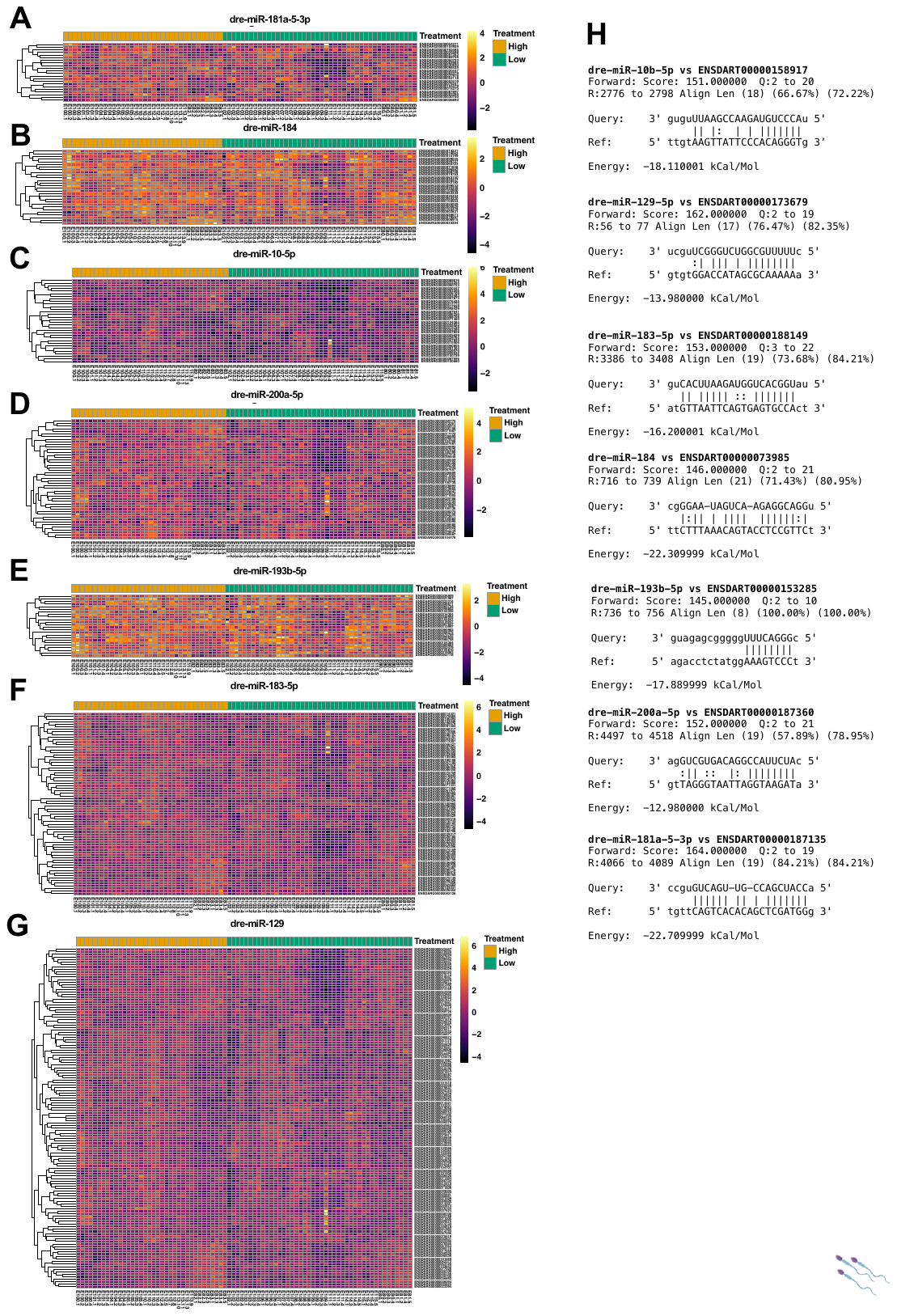

**Supplementary Figure 2.** Normalised expression of significantly differentially expressed genes targeted by the significantly differentially expressed miRNAs as predicted by miRanda, displaying log^2^ transformed CPM Z-score normalised counts. (A) dre-miR-181-5-3p, (B) dre-miR-184, (C) dre-miR-10b-5p, (D) dre-miR-200a-5p, (E) dre-miR-193b-5p, (F) dre-miR-183-5p, (G) dre-miR-129. (H) The 3’ UTR query sequence outputs from *miRanda* displaying predicted top-scoring miRNA target site.

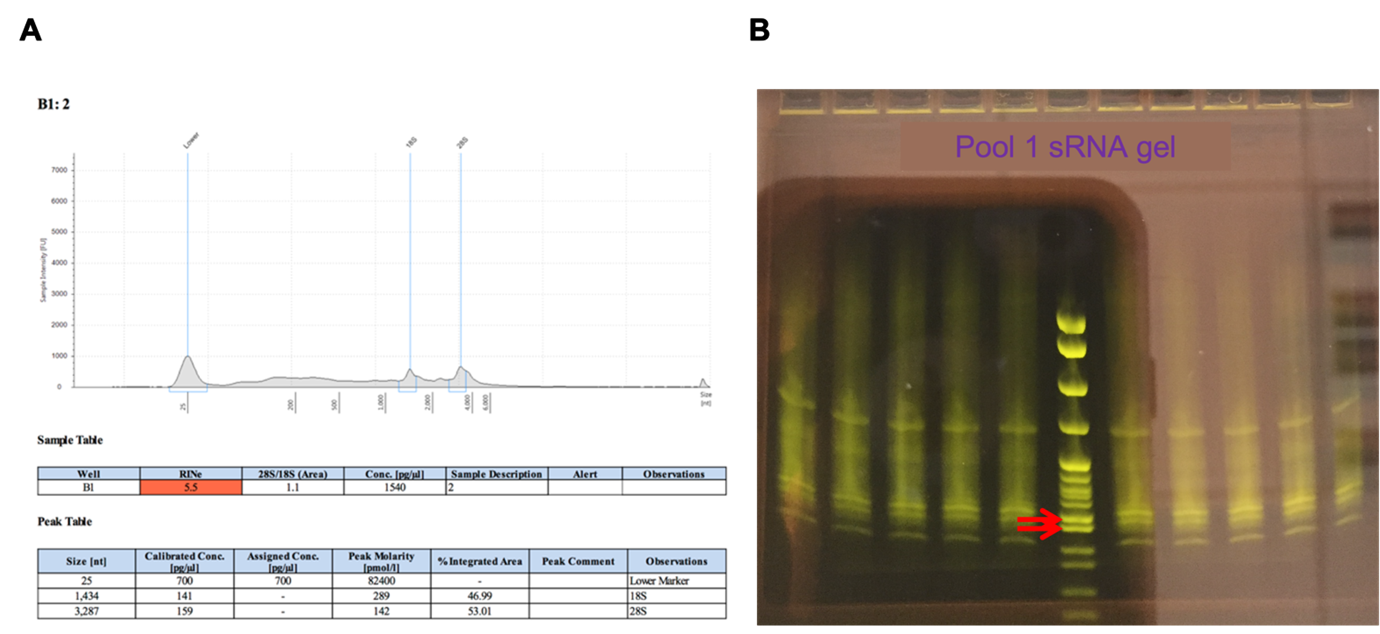

**Supplementary Figure 3.** Assessment of total RNA on TapeStation. (A) Samples show 18S and 28S peak. (B) A typical gel run showing the sRNA populations and the target regions that were cut from the gel, as indicated by the red arrows.

z

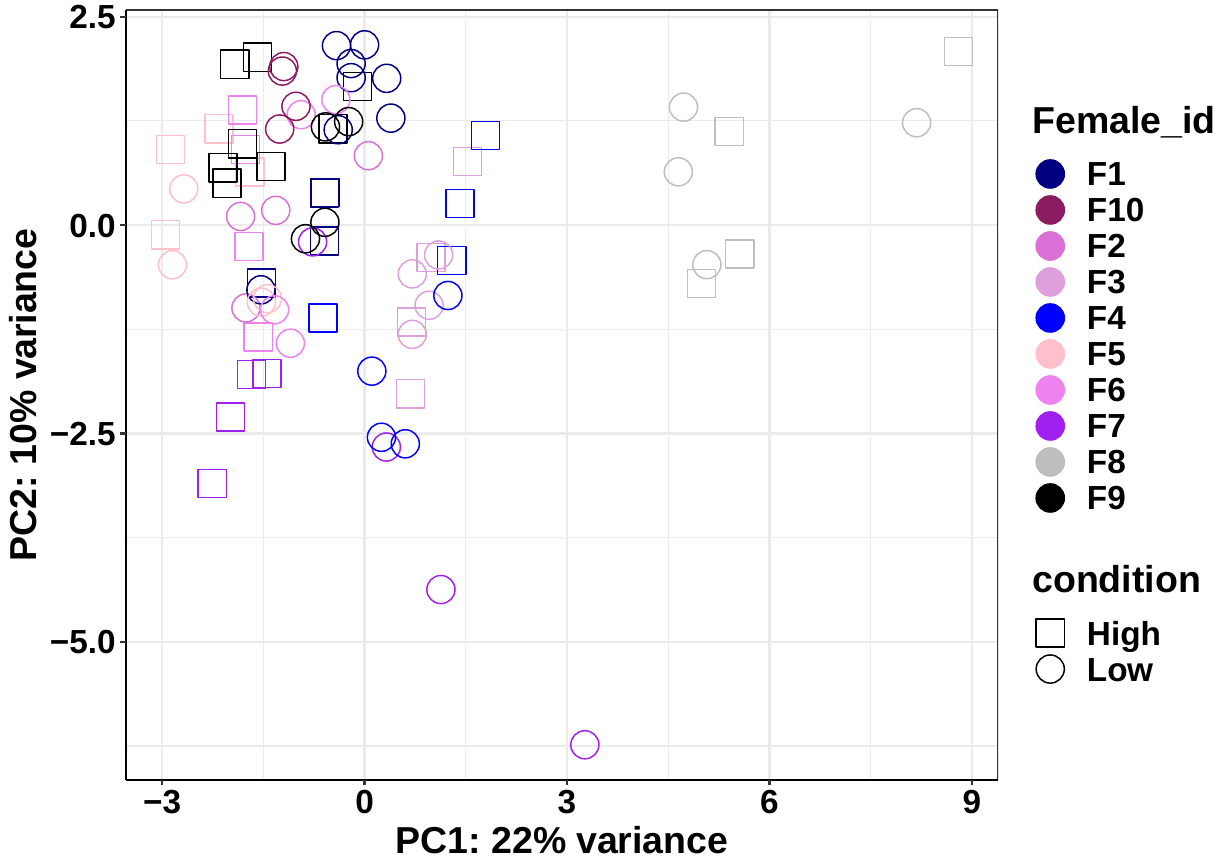

**A**

**C**

**B**

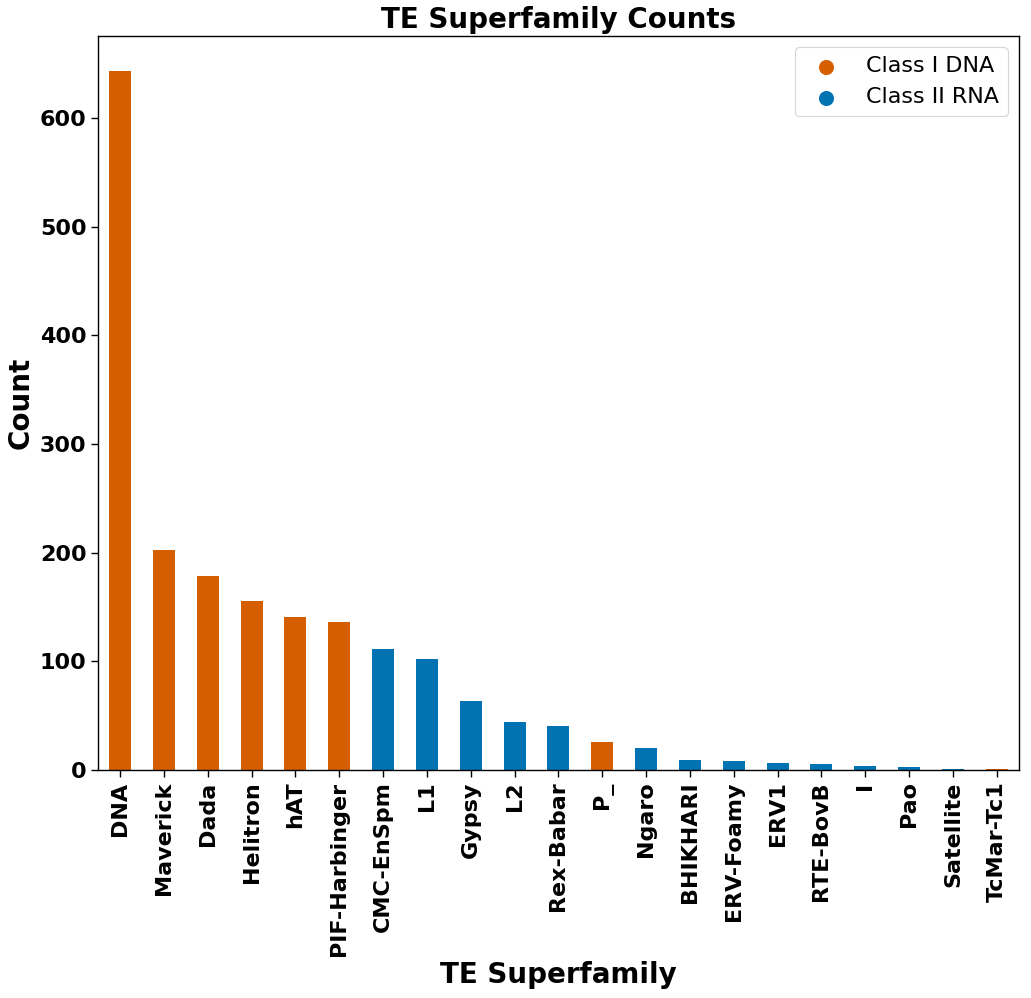

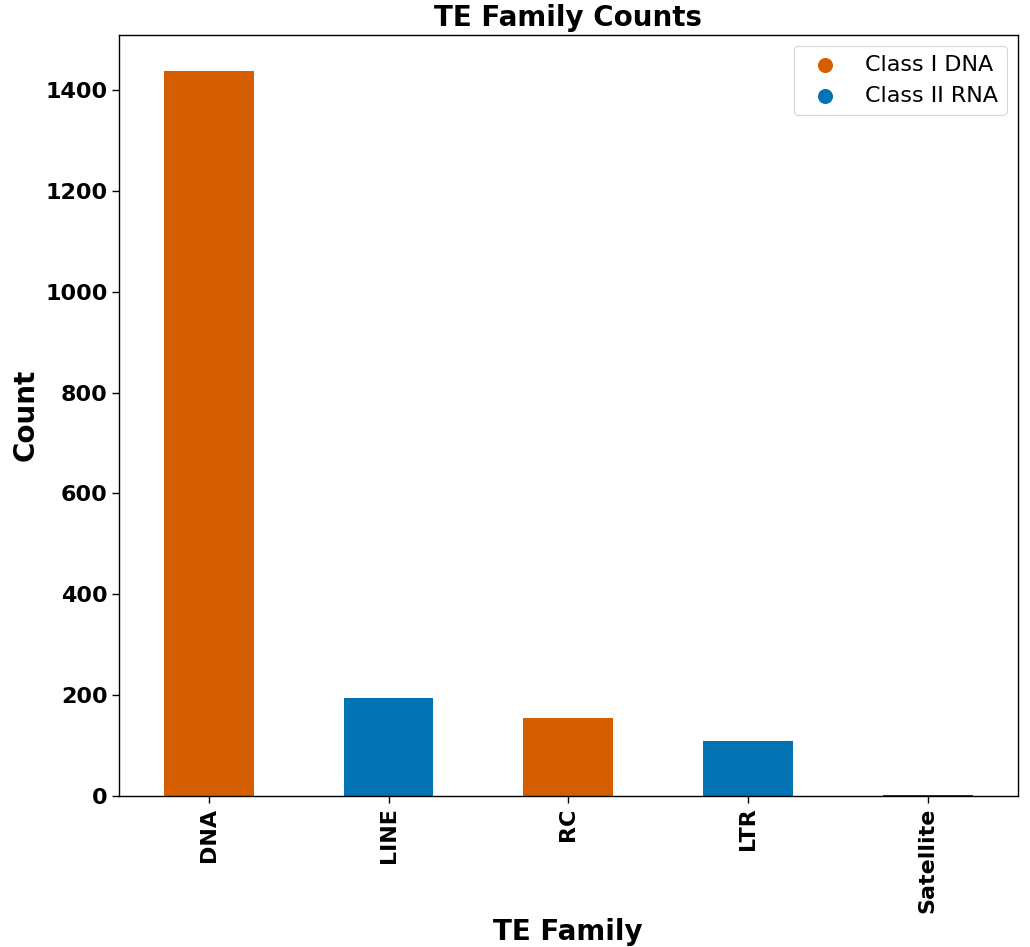

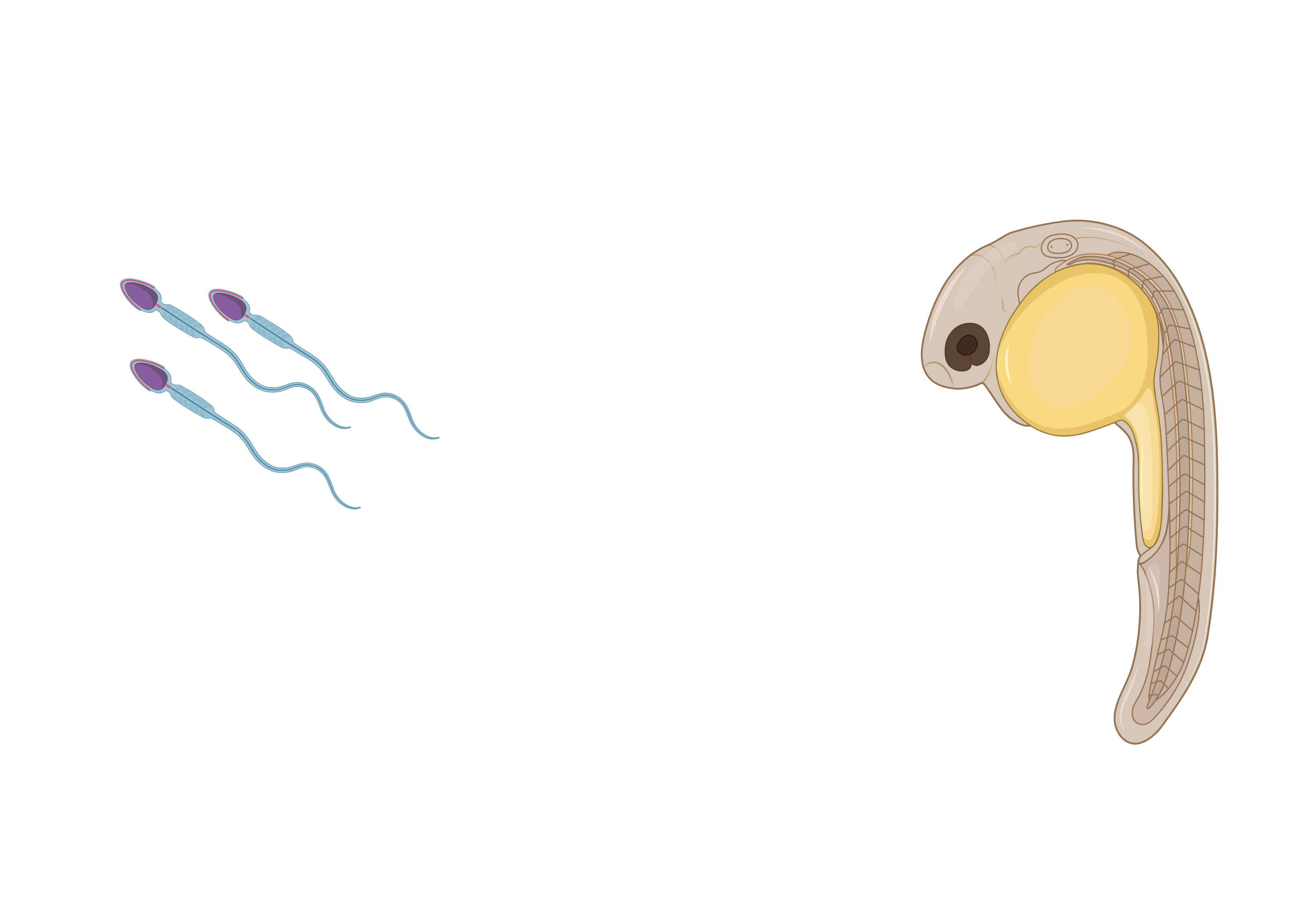

**Supplementary Figure 4.** TETranscripts clustering analysis and overlapping significantly differentially expressed TE families overlapping with genes count data **(A)** Principal Component Analysis with treatment (Low v High), and female ID as factors from DESeq2 on TETranscripts raw counts output. **(B)** Family counts from FishTEA pipleline for significantly differentially expressed TE’s that are overlapping with genes that are also significantly differentially expressed. Class I DNA transposons are displayed with orange bars, and Class II RNA retrotransposons are displayed with blue bars. **(C)** As with B, counts for TE superfamily.

**piR-dre-43599 complementary TEs**

**piR-dre-58396 complementary TEs**

**piR-dre-7058 complementary TEs**

**A**

**B**

**C**

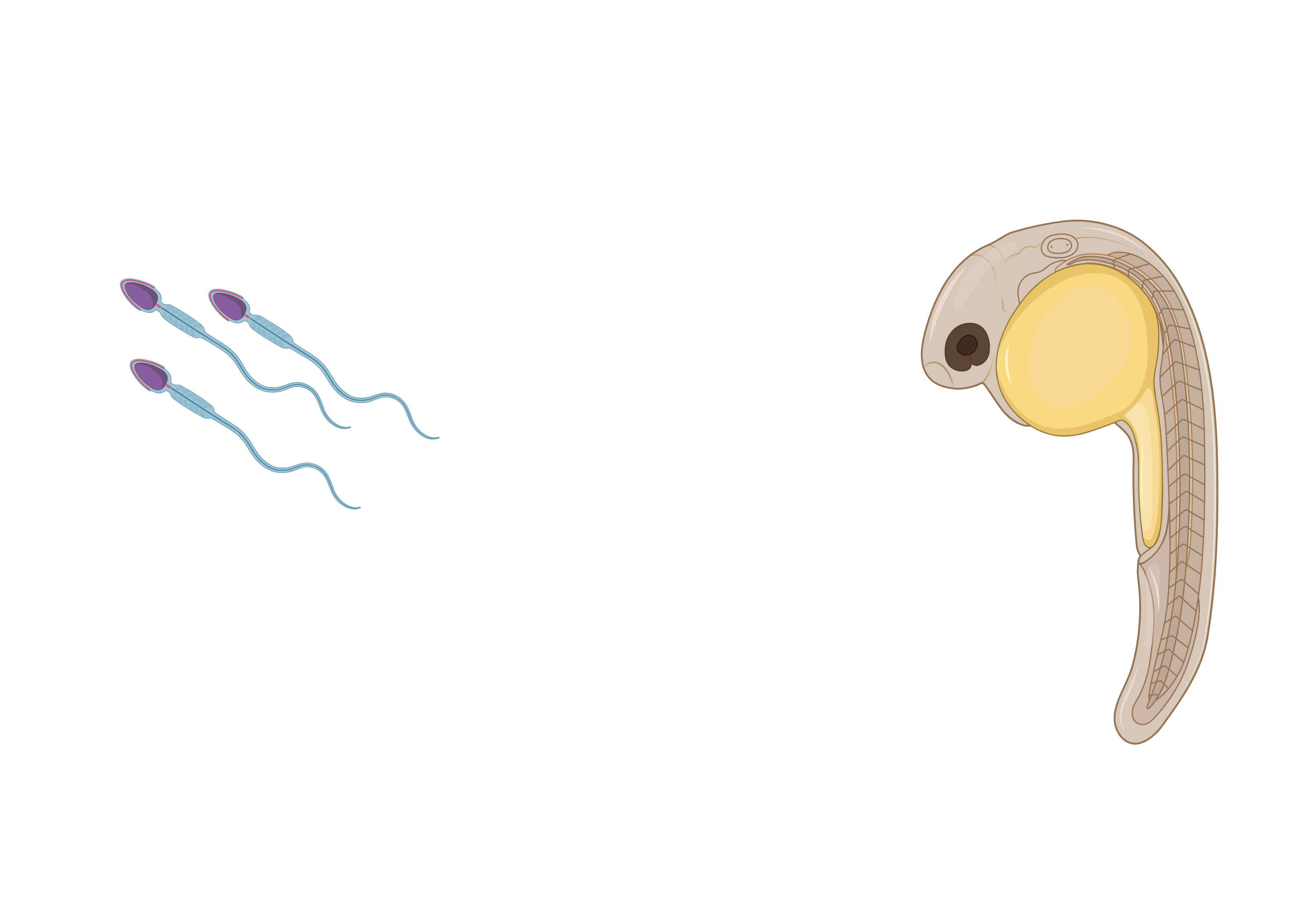

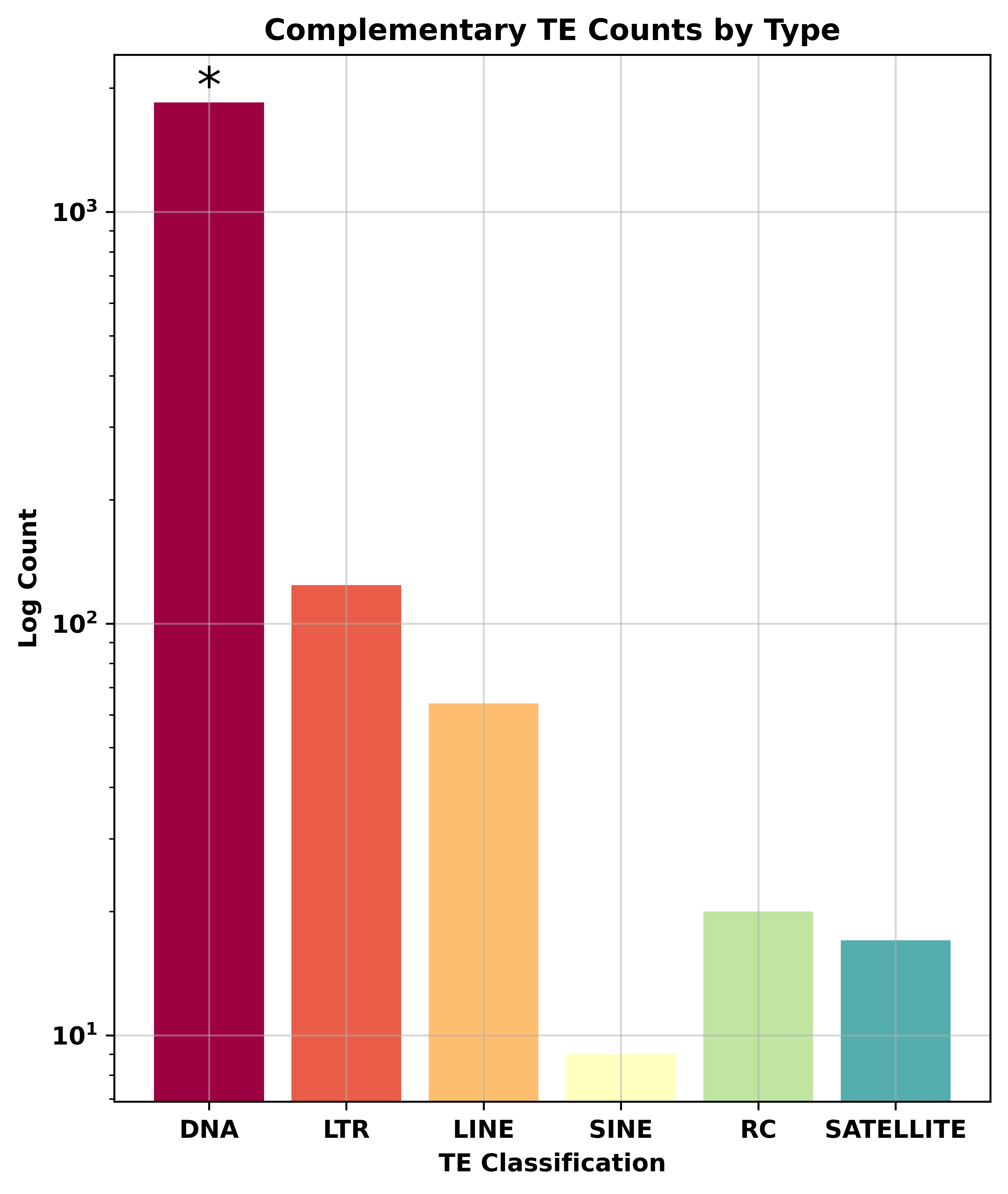

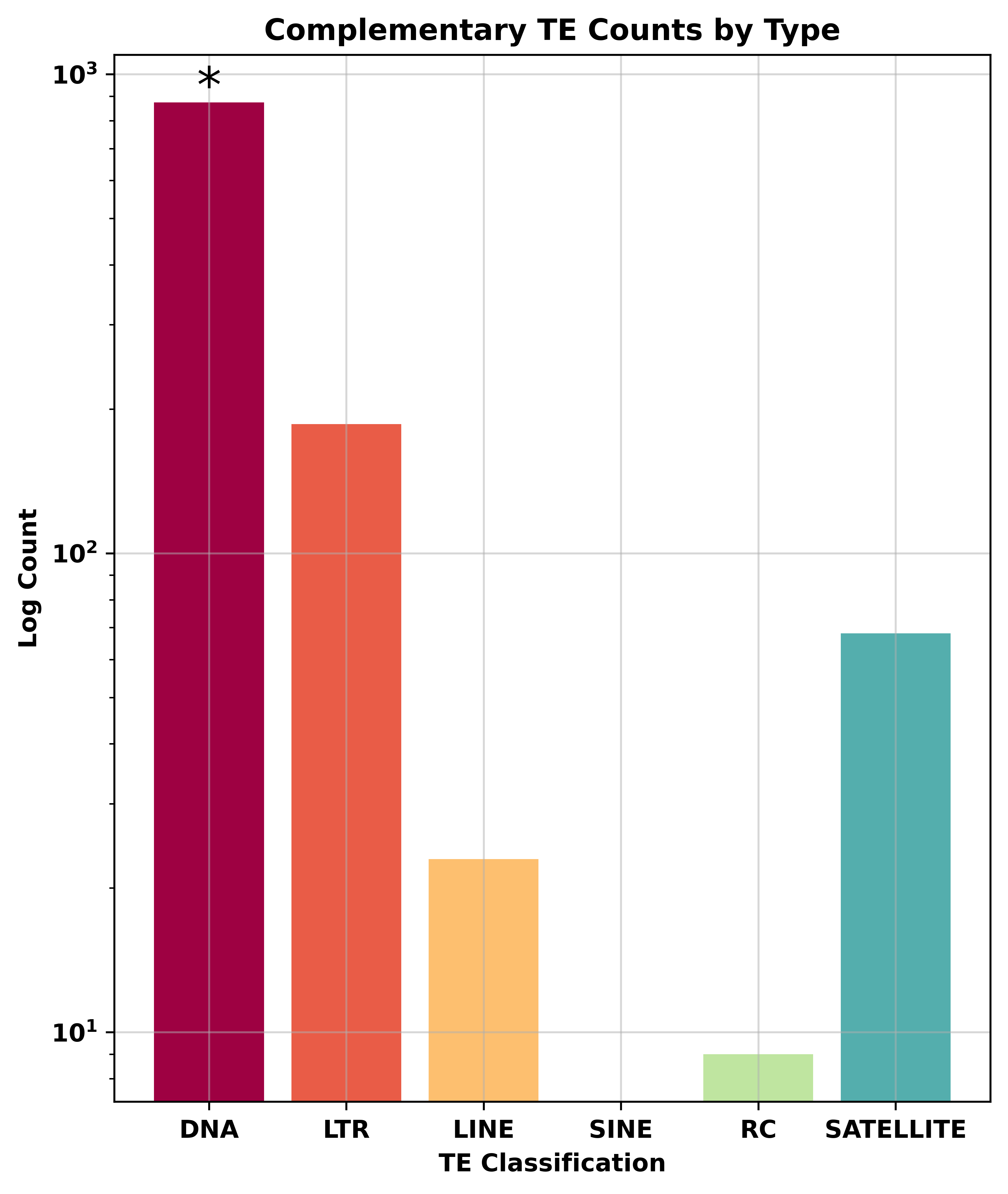

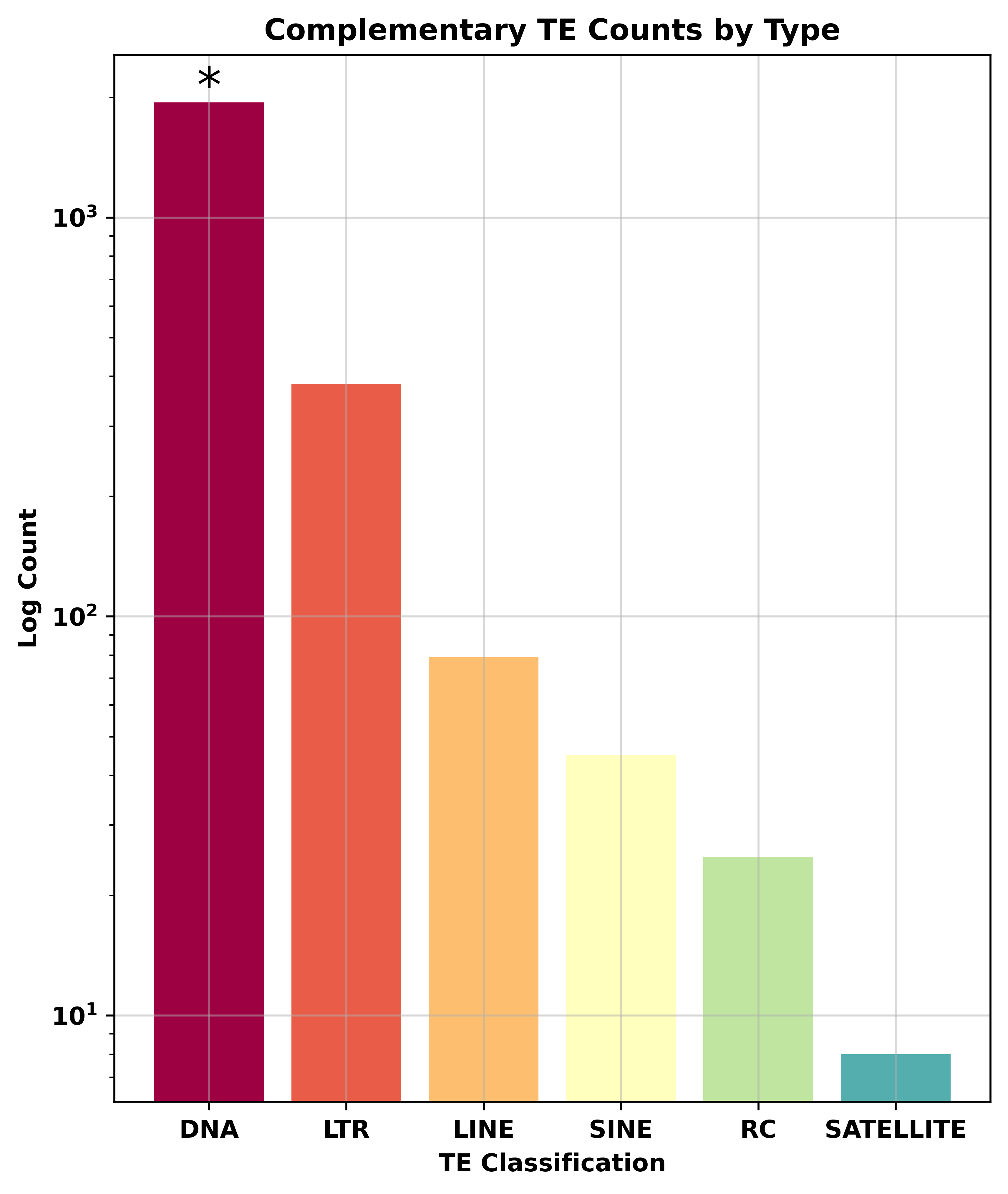

**Supplementary Figure 5.** PiRNA:TE complementarity analysis, results from FishPi. **(A)** Results for TE class complementary to piR-dre-43599, **(B)** piR-dre-7058, **(C)** and piR-dre-58396. *P* value “*” denotes *P* = < 0.001.

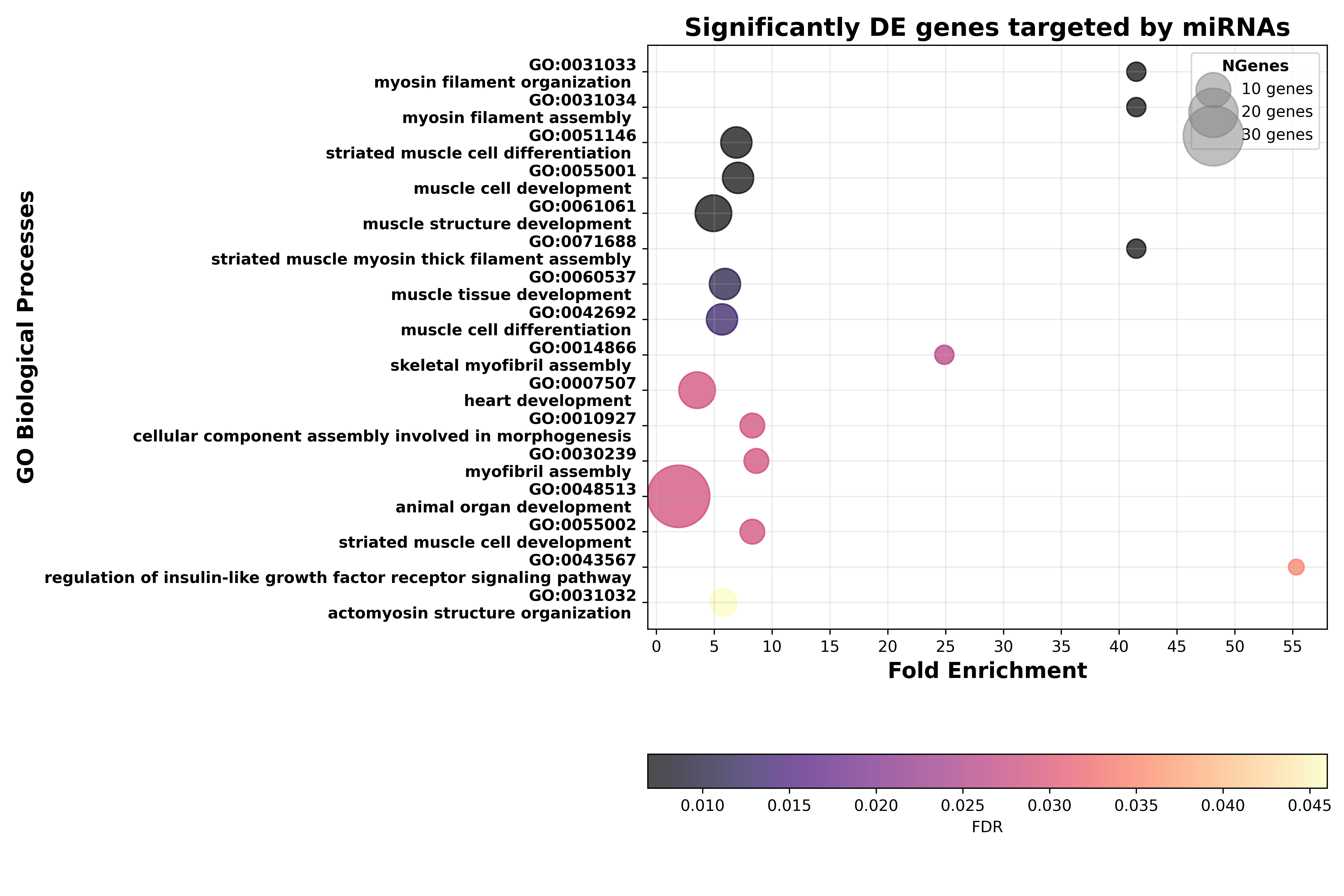

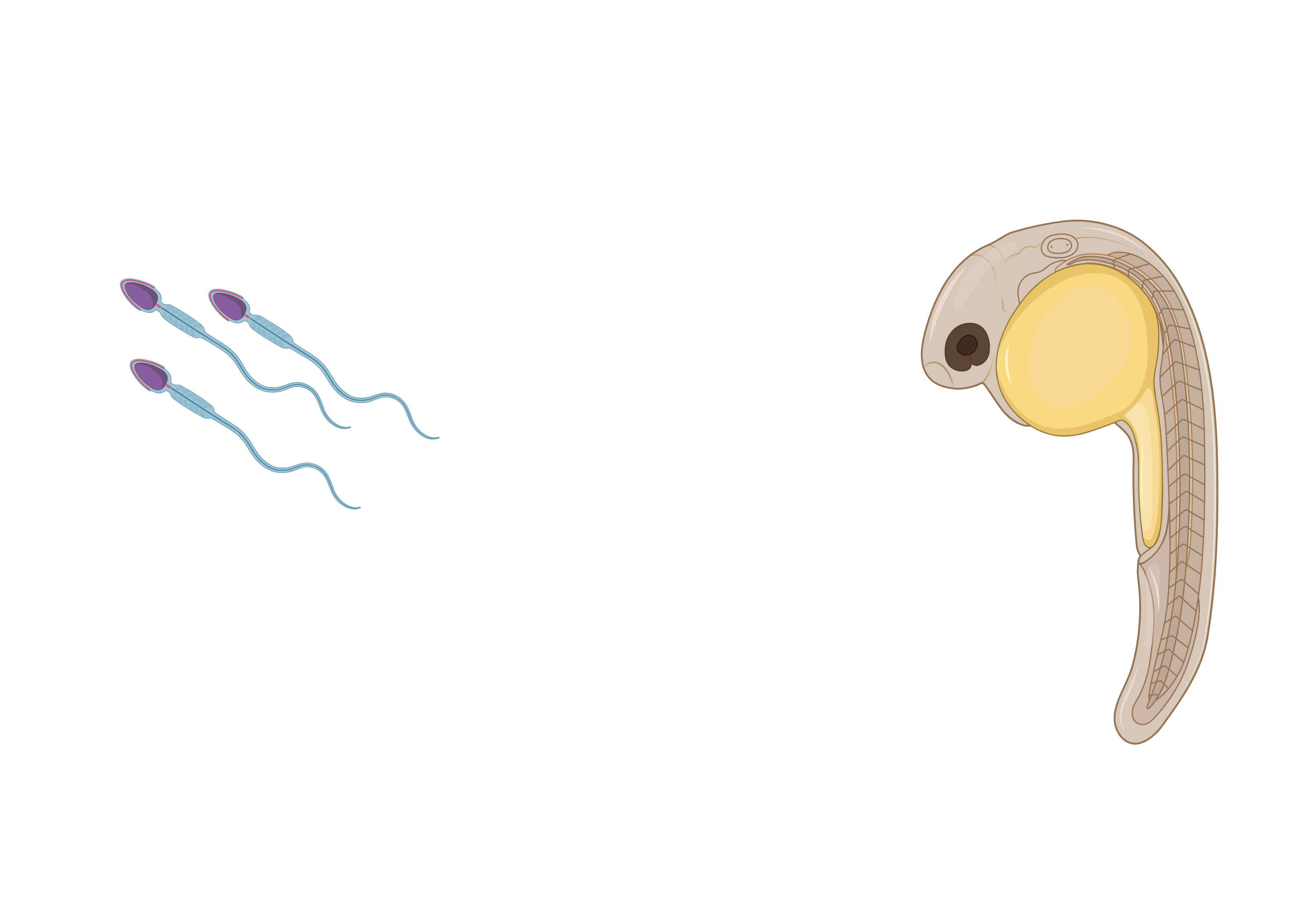

**Supplementary Figure 6.** sRNA-seq miRNA GO terms analysis. GO Biological processes on the significantly differentially expressed genes that are targets of the significantly differentially expressed miRNAs from miRanda analysis.

**Supplementary Tables**

**Supplementary Table S1.** Effects of social treatment and status on sperm VCL (N=86 males). Statistical parameters come from a linear mixed model (REML) with an Analysis of Deviance (Type III Wald chi-square tests). Male IDs were included as random variables. * indicate statistical significance at α<0.05.

|  | Chisq | Df | Pr(>Chisq) |
| --- | --- | --- | --- |
| (Intercept) | 5360.7945 | 1 | < 2.2e-16 * |
| Treatment | 0.0057 | 1 | 0.9395844 |
| Time | 5915.1316 | 1 | < 2.2e-16 * |
| Status | 0.0648 | 1 | 0.7990292 |
| Treatment:Time | 4.6236 | 1 | 0.0315359 * |
| Treatment:Time: Status | 13.9221 | 2 | 0.0009481 * |

**Supplementary Files**

**Supplementary materials, methods and figure legends**

resub_file_GoddenSilvaKiehl_Supplementary Material_heredity.docx

**Supplementary File S1- List of piRNA cluster full names in Fig. 4B**

Supp_File_1_piRNAs_heatmap_fullnames.xlsx

**Supplementary File S2 – Phenotypic data**

Supp_File_6_sperm_motility_data.csv

**Supplementary File S3 & S4- raw counts and metadata file RNA seq**

Supp_File_3_RNA_seq_rawcounts.csv

Supp_File_4_rnaseq_metadata.csv

**Supplementary File S5 & S6- raw counts and metadata file sRNA-seq: miRNA data**

Supp_File_5_40_mirna_counts_LF.xlsx

Supp_File_6_40_mirna_metadata_LF.csv

**Supplementary File S7 & S8- raw counts and metadata file sRNA-seq: piRNA data**

Suppl_File_7_piRNA_raw_counts.csv

Supp_File_8_pirna_metadata.csv

**Supplementary File S9 & S10- TEtranscripts RNAseq raw counts output and metadata file**

Suppl_File_9_tetranscripts_raw_counts_Teonly.xlsx

Suppl_File_10_metadata_rnaseq_TEtranscripts.csv
